# A highly conserved and globally prevalent cryptic plasmid is among the most numerous mobile genetic elements in the human gut

**DOI:** 10.1101/2023.03.25.534219

**Authors:** Emily C Fogarty, Matthew S Schechter, Karen Lolans, Madeline L. Sheahan, Iva Veseli, Ryan Moore, Evan Kiefl, Thomas Moody, Phoebe A Rice, Michael K Yu, Mark Mimee, Eugene B Chang, Sandra L Mclellan, Amy D Willis, Laurie E Comstock, A Murat Eren

**Affiliations:** Committee on Microbiology, University of Chicago, Chicago, IL 60637, USA; Duchossois Family Institute, University of Chicago, Chicago, IL 60637, USA; Department of Medicine, University of Chicago, Chicago, IL 60637, USA; Department of Microbiology, University of Chicago, Chicago, IL, 60637, USA; Graduate Program in Biophysical Sciences, University of Chicago, Chicago, IL 60637, USA; Center for Bioinformatics and Computational Biology, University of Delaware, Newark, DE, USA; Department of Systems Biology, Columbia University, New York, NY, 10032 USA; Department of Biochemistry, University of Chicago, Chicago, IL, 60637, USA; Toyota Technological Institute at Chicago; Pritzker School of Molecular Engineering, The University of Chicago, Chicago, IL 60637, USA; School of Freshwater Sciences, University of Wisconsin-Milwaukee, Milwaukee, WI, 53204, USA; Department of Biostatistics, University of Washington, Seattle, WA, 98195, USA; Marine Biological Laboratory, Woods Hole, MA, 02543, USA; Alfred Wegener Institute, Helmholtz Centre for Polar and Marine Research, 27570 Bremerhaven, Germany; Institute for Chemistry and Biology of the Marine Environment, University of Oldenburg, 26129 Oldenburg, Germany; Helmholtz Institute for Functional Marine Biodiversity, 26129 Oldenburg, Germany

## Abstract

Plasmids are extrachromosomal genetic elements that often encode fitness enhancing features. However, many bacteria carry ‘cryptic’ plasmids that do not confer clear beneficial functions. We identified one such cryptic plasmid, pBI143, which is ubiquitous across industrialized gut microbiomes, and is 14 times as numerous as crAssphage, currently established as the most abundant genetic element in the human gut. The majority of mutations in pBI143 accumulate in specific positions across thousands of metagenomes, indicating strong purifying selection. pBI143 is monoclonal in most individuals, likely due to the priority effect of the version first acquired, often from one’s mother. pBI143 can transfer between Bacteroidales and although it does not appear to impact bacterial host fitness *in vivo*, can transiently acquire additional genetic content. We identified important practical applications of pBI143, including its use in identifying human fecal contamination and its potential as an inexpensive alternative for detecting human colonic inflammatory states.

## INTRODUCTION

The tremendous density of microbes in the human gut provides a playground for the contact-dependent transfer of mobile genetic elements ^1^ including plasmids. Plasmids are typically defined as extrachromosomal elements that replicate autonomously from the host chromosome ^1–4^. In addition to being a workhorse for molecular biology, plasmids have been extensively studied for their ability to expedite microbial evolution ^5^ and enhance host fitness by providing properties such as antibiotic resistance, heavy metal resistance, virulence factors, or metabolic functions ^6–11^.

Plasmids have been a major focus of microbiology not only for their biotechnological applications to molecular biology ^12–15^ but also for their role in the evolution and dissemination of genes for antibiotic resistance ^16, 17^, which is a growing global public health concern ^18^. However, outside the spotlight lie a group of plasmids that appear to lack genetic functions of interest and that do not contain genes encoding obvious beneficial functions to their hosts ^19, 20^. Such ‘cryptic plasmids’ are typically small and multi-copy ^21^, and are often difficult to study as they lack any measurable phenotypes or selectable markers ^22, 23^, despite their presence in a broad range of microbial taxa ^24–27^. In the absence of a clear advantage to their hosts, and the presumably non-zero cost of their maintenance, these plasmids are often described as selfish elements ^28^ or genetic parasites ^29^. While they may provide unknown benefits to their hosts, a high transfer rate could also be a factor that enables cryptic plasmids to counteract the negative selection pressure of their maintenance ^29–31^.

Analyses of cryptic plasmids are often performed on monocultured bacteria, limiting insights into the ecology of cryptic plasmids in their host’s natural environment. However, recent advances in shotgun metagenomics ^30^ and *de novo* plasmid prediction algorithms ^31–40^ offer a powerful means to bridge this gap. For instance, in a recent study we characterized over 68,000 plasmids from the human gut ^40^ and observed that the most prevalent known plasmid across geographically diverse human populations was a cryptic plasmid, called pBI143. Here we conduct an in-depth characterization of this cryptic plasmid through ’omics and experimental approaches to study its genetic diversity, host range, transmission routes, impact on the bacterial host, and associations with health and disease states. Our findings reveal the astonishing success of pBI143 in the human gut, where it occurs in up to 92% of individuals in industrialized countries with copy numbers 14 times higher on average than crAssphage, the most abundant phage in the human gut. We also demonstrate the potential of pBI143 as a cost-effective biomarker to assess the extent of stress that microbes experience in the human gut, and as a sensitive means to quantify the level of human fecal contamination in environmental samples.

## RESULTS

### pBI143 is extremely prevalent across industrialized human gut microbiomes

pBI143 (accession ID U30316.1) is a 2,747 bp circular plasmid first identified in 1985 ^41^ in *Bacteroides fragilis* ^42^, an important member of the human gut microbiome that is frequently implicated in states of health ^43–45^ and disease ^46, 47^. pBI143 encodes only two annotated genes: a mobilization protein (*mobA*) and a replication protein (*repA*) (Fig. 1A). Due to the desirable features for cloning such as a high copy number and genetic stability, pBI143 has been primarily used as a component of *E. coli-Bacteroides* shuttle vectors ^42^. The absence of any ecological studies of pBI143 prompted us to characterize it further beginning with a characterization of its genetic diversity.

**Fig. 1.**
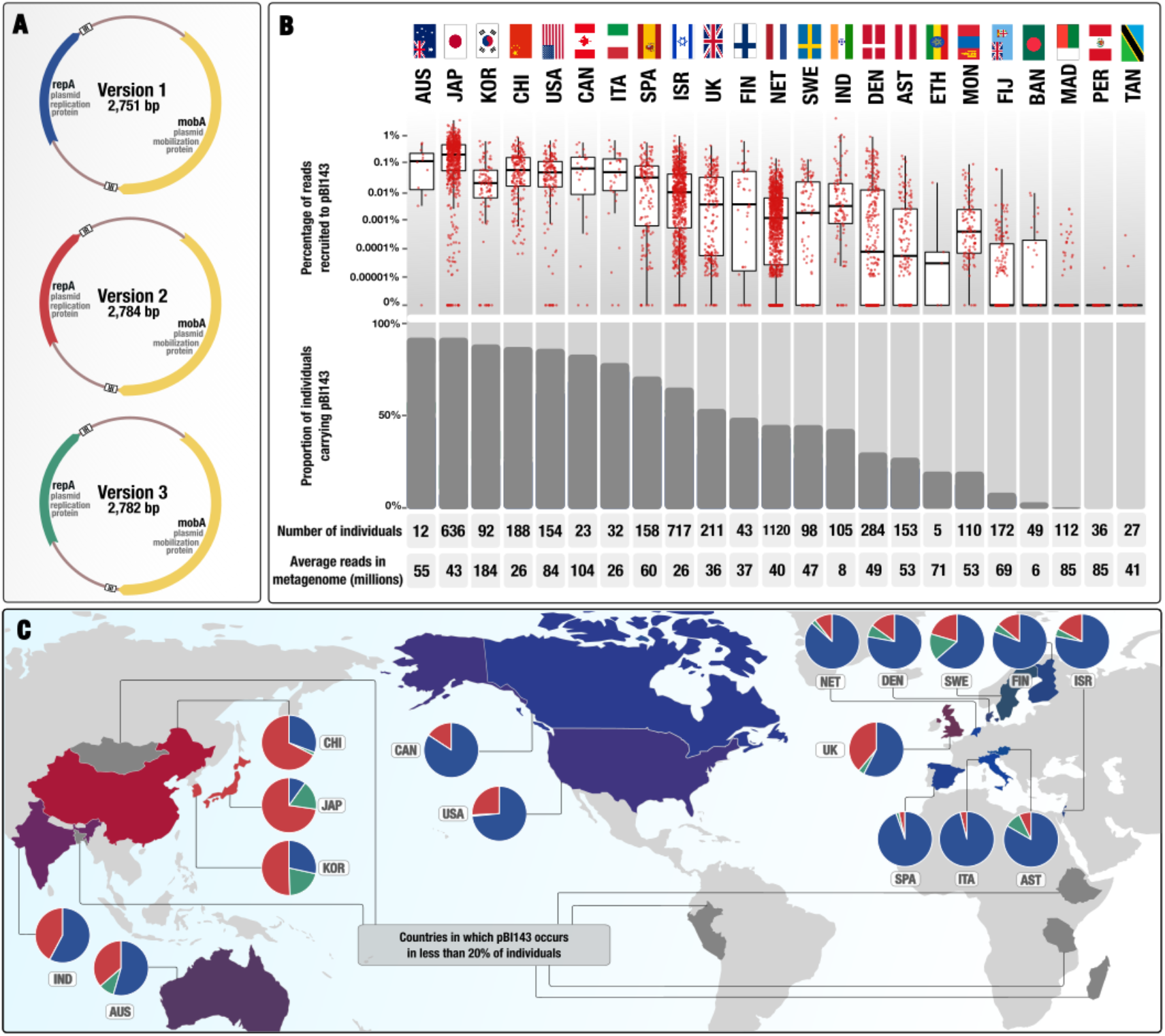
pBI143 prevalence and abundance in globally distributed human populations. (A) Plasmid maps of the three distinct versions of pBI143, which differ primarily in the *repA* gene. IR = inverted repeat. The *repA* genes are colored according to Version 1 (blue), Version 2 (red) and Version 3 (green). (B) Read recruitment results from 4,516 metagenomes originating from 23 globally representative countries and mapped to pBI143. Top: The percentage of reads in each metagenome that mapped to pBI143 normalized by number of reads in the metagenome. Bottom: The proportion of individuals in a country that have pBI143 in their gut. Each red dot represents an individual metagenome. (C) Countries that are represented in our collection of 4,516 global adult gut metagenomes. Each country’s pie chart is colored based on the version(s) of pBI143 that is most prevalent in that country (Version 1 = blue, Version 2 = red, Version 3 = green). Each country is colored based on the proportion of Version 1, 2 or 3 present in the population, or gray if fewer than 20% of individuals carry pBI143. Pie charts show the proportions of pBI143 versions in all individuals that carry it within a country.

To comprehensively sample the diversity of pBI143, we screened 2,137 individually assembled human gut metagenomes (Supplementary Table 1) for pBI143-like sequences. By surveying all contigs using the known pBI143 sequence as reference, we found three distinct versions of pBI143 (Fig. 1A), all of which had over 95% nucleotide sequence identity to one another throughout their entire length except at the *repA* gene, where the sequence identity was as low as 75% with a maximum of 81% between Version 1 and Version 2 (Supplementary Table 2).

We then sought to quantify the prevalence of pBI143 across global human populations using a metagenomic read recruitment survey with an expanded set of 4,516 publicly available gut metagenomes from 23 countries ^48,49,50,51,52,53,54,55,56,57,58,59,60,61,62,63,55,64,65,66,67,68,66,69,70^ (Supplementary Table 1). Recruiting metagenomic short reads from each gut metagenome using each pBI143 version independently (Supplementary Fig. 1, Supplementary Table 3), we found that pBI143 was present in 3,295 metagenomes, or 73% of all samples (Fig. 1B, see Methods for the ‘detection’ criteria). However, the prevalence of pBI143 was not uniform across the globe (Fig. 1B): pBI143 occurred predominantly in metagenomes of individuals who lived in relatively industrialized countries, such as Japan (92% of 636 individuals) and the United States (86% of 154 individuals). We rarely detected pBI143 in individuals who lived in relatively non-industrialized countries such as Madagascar (0.8% of 112 individuals) or Fiji (8.7% of 172 individuals). This differential coverage is likely due to the non-uniform distribution of *Bacteroides* populations, which tend to dominate individuals who live in relatively more industrialized countries ^71^. Within each individual, pBI143 was often highly abundant (Fig. 1B), and despite its small size, it often recruited 0.1% to 3.5% of all metagenomic reads with a median coverage of over 7,000X (Supplementary Fig. 1, Supplementary Table 3). In one extreme example, pBI143 comprised an astonishing 7.5% of all reads in an infant gut metagenome from Italy, with a metagenomic read coverage exceeding 54,000X (Supplementary Table 3).

The distribution of pBI143 versions across human populations was also not uniform as different versions of pBI143 tended to be dominant in different geographic regions. pBI143 Version 1 (98% identical to the original reference sequence for pBI143 ^41^) dominated individuals in North America and Europe, and occurred on average in 82.5% of all samples that carry pBI143 from Austria, Canada, Denmark, England, Finland, Italy, Netherlands, Spain, Sweden and the USA (Fig. 1C, Supplementary Table 3). In contrast, pBI143 Version 2 dominated countries in Asia and occurred in 63.6% of all samples that carry pBI143 in China, Japan, and Korea (Fig. 1C, Supplementary Table 3). pBI143 Version 3 was relatively rare, comprising only 7.4% of pBI143-positive samples, and mostly occurred in individuals from Japan, Korea, Australia, Sweden, and Israel (Fig. 1C, Supplementary Table 3).

The extremely high prevalence and coverage of pBI143 suggests that it is likely one of the most numerous genetic elements in the gut microbiota of individuals from industrialized countries. We compared the prevalence and relative abundance of pBI143 to crAssphage, a 97 kbp bacterial virus that is widely recognized as the most abundant family of viruses in the human gut ^72^. pBI143 was more prevalent (73% vs 27%) in our metagenomes than crAssphage, although individual samples differed widely with respect to the abundance of these two elements in a given individual (Supplementary Table 3). The average percentage of metagenomic reads recruited by pBI143 and crAssphage were 0.05% and 0.13%, respectively. However, taking into consideration that crAssphage is approximately 36 times larger than pBI143, and assuming that average coverage is an acceptable proxy to the abundance of genetic entities, these data suggest that on average pBI143 is 14 times more numerous than crAssphage in the human gut.

Overall, these data demonstrate that pBI143 is one of the most widely distributed and numerous genetic elements in the gut microbiomes of industrialized human populations world-wide.

### pBI143 is specific to the human gut and hosted by a wide range of Bacteroidales species

Interestingly, the detection patterns of pBI143 in metagenomes differed from the detection patterns we observed for its *de facto* host *Bacteroides fragilis* in the same samples; *B. fragilis* and pBI143 co-occurred in only 41% of the metagenomes. Sequencing depth did not explain this observation, as pBI143 was highly covered (i.e., >50X) in 25% of metagenomes where *B. fragilis* appeared to be absent (Supplementary Table 11), suggesting that the host range of pBI143 extends beyond *B. fragilis*.

To investigate the host range of pBI143, we employed a collection of bacterial isolates from the human gut, which contained 717 genomes that represented 104 species in 54 genera (Supplementary Table 4). We found pBI143 in a total of 82 isolates that resolved to 11 species across 3 genera: *Bacteroides*, *Phocaeicola*, and *Parabacteroides*. Many of the pBI143-carrying isolates of distinct species were from the same individuals, suggesting that pBI143 can be mobilized between species. To confirm this, we inserted a tetracycline resistance gene, *tetQ*, into pBI143 in the *Phocaeicola vulgatus* isolate MSK 17.67 (Supplementary Fig. 2, Supplementary Table 4) and tested the ability of this engineered pBI143 to transfer to two strains of two different families of Bacteroidales, *Bacteroides ovatus* D2 and *Parabacteroides johnsonii* CL02T12C29. In these assays, we found that pBI143 was indeed transferred from the donor to the recipient strains at a frequency of 5 x 10^-7^ and 3 x 10^-6^transconjugants per recipient, respectively (Supplementary Fig. 2).

Given the broad host range of pBI143, one interesting question is whether the ecological niche boundaries of pBI143 hosts exceed a single biome, since the members of the order Bacteroidales are not specific to the human gut and do occur in a wide range of other habitats from non-human primate guts ^73^ to marine systems ^74^. To investigate whether pBI143 might exist in non-human environments, we searched for pBI143 in metagenomes from coastal and open ocean samples ^75, 76^, captive macaques ^73^, human-associated pets ^77^, and sewage samples from across the globe ^78^. The plasmid was absent from all non-human associated samples, but as expected, was present in sewage (Supplementary Fig. 3, Supplementary Table 3, Supplementary Text). Given the absence of pBI143 in non-human associated habitats, we also screened metagenomes from human skin and oral cavity ^70^. Unlike the extremely high presence of pBI143 in the human gut, pBI143 was poorly detected both in samples from skin and the oral cavity (Supplementary Text). Finally, we designed and tested a highly specific qPCR assay for pBI143 (Supplementary Table 5) to confirm its specificity to the human gut. While there was a robust amplification of pBI143 from sewage samples confirming our insights from metagenomic coverages (Fig. 2), pBI143 was virtually absent in dog, alligator, raccoon, horse, pig, deer, cow, chicken, goose, cat, rabbit, deer, or gull fecal samples (Supplementary Table 6). The only exception was the relatively low copy number (i.e., 73-fold less than human fecal content of sewage) in three of the four cats tested.

**Fig. 2.**
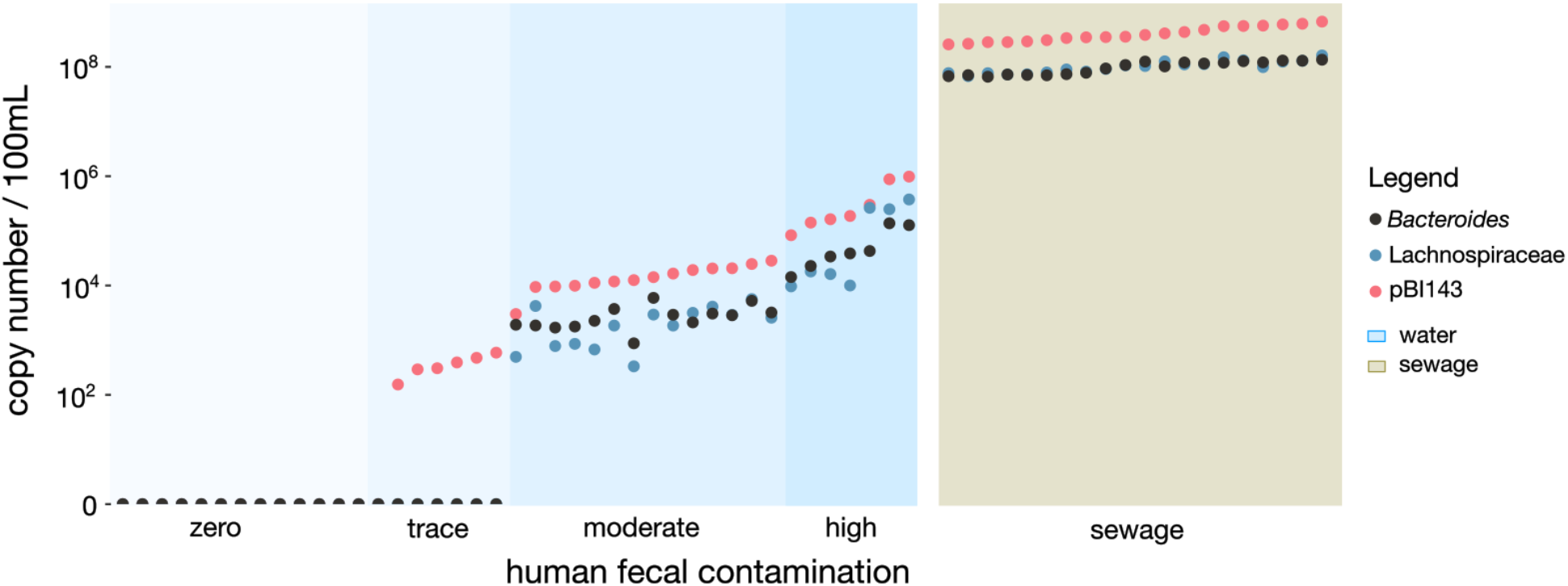
Detection of pBI143 and two established human fecal markers in water and sewage samples. Copy number of pBI143, human *Bacteroides* or Lachnospiraceae as measured by qPCR. Zero, trace, moderate, high and sewage categories and sample order designations are determined based on pBI143 copy number. Trace indicates one of the established markers was detected but was below the level of quantification. The blue background indicates water samples and the beige background indicates samples from sewage.

The near-absolute exclusivity of pBI143 to the human gut presents practical opportunities, such as the accurate detection of human fecal contamination outside the human gut. Using the same PCR primers, we also amplified pBI143 from water and sewage samples and compared its sensitivity to the gold standard markers currently used for detecting human fecal contamination in the environment (16S rRNA gene amplification of human *Bacteroides* and Lachnospiraceae) ^79, 80^. pBI143 had higher amplification in all 41 samples where *Bacteroides* and Lachnospiraceae were also detected (Fig. 2). pBI143 was also amplified in 6 samples with no *Bacteroides* or Lachnospiraceae amplification, suggesting it is a highly sensitive marker for detecting the presence of human-specific fecal material.

Overall, these data show that pBI143 has a broad range of Bacteroidales species, is highly specific to the human gut environment, and can serve as a sensitive biomarker to detect human fecal contamination.

### pBI143 is monoclonal within individuals, and its variants across individuals are maintained by strong purifying selection

So far, our investigation of pBI143 has focused on its ecology. Next, we sought to understand the evolutionary forces that have conserved the pBI143 sequence by quantifying the sequence variation among the three distinct versions and examining the distribution of single nucleotide variants (SNVs) within and across globally distributed individuals. Across the three versions, both pBI143 genes had low dN/dS values (*mobA* = 0.11, *repA* = 0.04), suggesting the presence of strong forces of purifying selection acting on *mobA* and *repA* resulting in primarily synonymous substitutions. While the comparison of the three representative sequences provide some insights into the conserved nature of pBI143, it is unlikely they capture its entire genetic diversity across gut metagenomes.

To explore the pBI143 variation landscape, we analyzed metagenomic reads that matched the Version 1 of *mobA* to gain insights into the population genetics of pBI143 in naturally occurring habitats through single-nucleotide variants (SNVs). Since the *mobA* gene was more conserved across distinct versions of the plasmid compared to the *repA* gene, focusing on *mobA* enabled characterization of variation from all plasmid versions using a single read recruitment analysis. Surprisingly, the vast majority (83.2%) of the nucleotide positions that varied in any metagenome matched a nucleotide position that was variable between at least one pair of the three plasmid versions (Fig. 3A, Supplementary Table 7). In other words, pBI143 variation across metagenomes was predominantly localized to certain nucleotide positions that differed between the representative sequences of pBI143 for Version 1, 2 and 3, indicating that the three representative versions capture the majority of permissible pBI143 variation within our collection of gut metagenomes. Indeed, only 24.5% of metagenomes had more than three novel SNVs that were not present in at least one plasmid version, and 84.8% of metagenomes had pBI143 sequences that were within 2-nucleotide distance of one of the three versions. In addition to the primarily localized variation of pBI143, we also observed that the vast majority of SNVs were fixed within a metagenome (i.e., a ‘departure from consensus’ value of ∼0, see Methods), suggesting that most humans carry a monoclonal population of pBI143 with little to no within-individual variation (Fig. 3C, Supplementary Table 7).

**Fig. 3.**
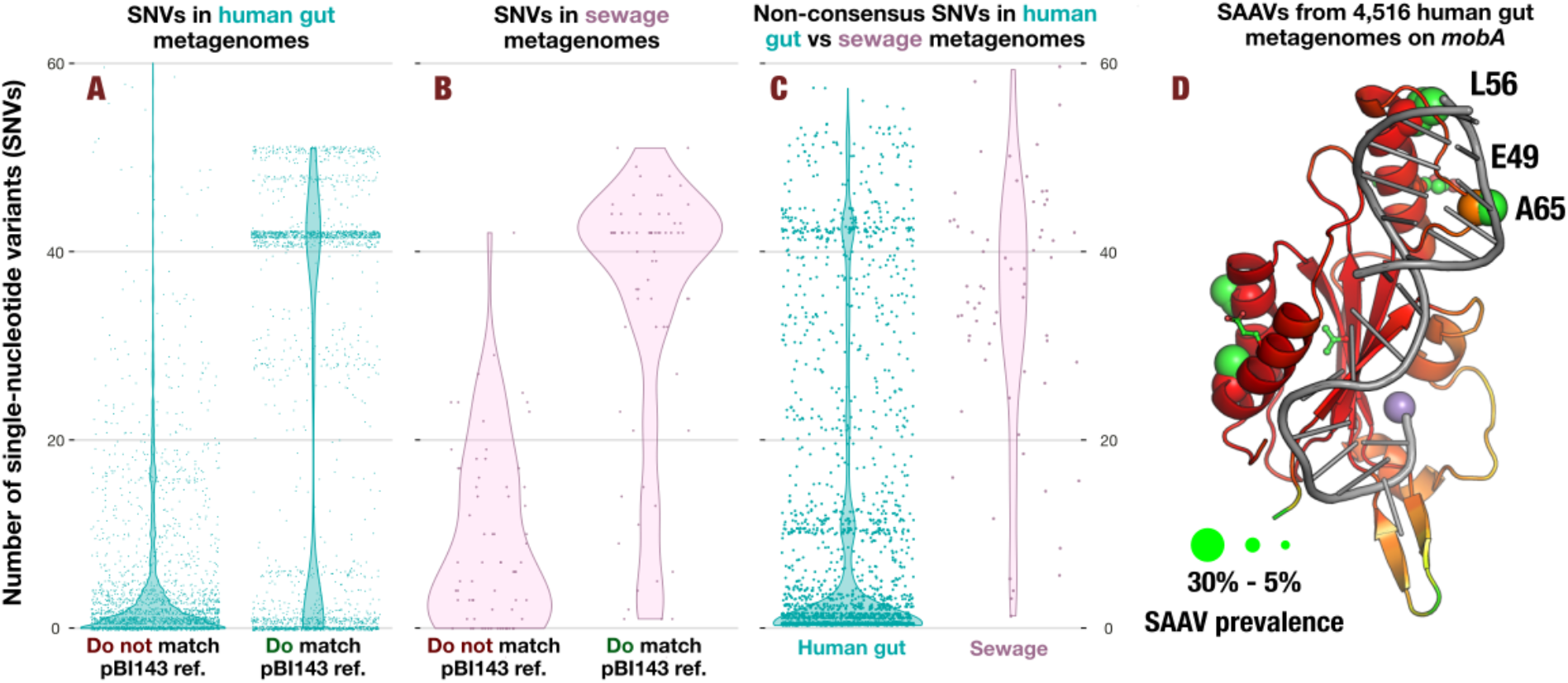
The mutational landscape of pBI143 in sewage and the human gut. **(A)** The proportion of SNVs across 4,516 human gut metagenomes that are present in the same location (match) or different locations (do not match) as variation in one of the versions of pBI143 (turquoise). Each point is a single metagenome. **(B)** The proportion of SNVs across 68 sewage gut metagenomes that are present in the same location (match) or different locations (do not match) as variation in one of the versions of pBI143 (pink). **(C)** Non-consensus SNVs present in 4,516 human gut metagenomes and 68 sewage metagenomes. **(D)** AlphaFold 2 predicted structure of the catalytic domain of MobA with single amino acid variants from all 4,516 human gut metagenomes superimposed as ball-and-stick residues. oriT DNA (gray) and a Mn2+ ion marking the active site (purple) were modeled based on 4lvi.pdb ^81^. The size of the ball-and-stick spheres indicate the proportion of samples carrying variation in that position (the larger the sphere, the more prevalent the variation at the residue) and the color is in CPK format. The color of the ribbon diagram indicates the pLDDT from AlphaFold 2 with red = very high (> 90 pLDDT) and orange = confident (80 pLDDT).

Next, we sought to investigate the functional context of non-synonymous environmental variants of MobA given its structure. For this, we employed single-amino acid variants ^82^ (SAAVs) we recovered from gut metagenomes and superimposed them on the AlphaFold 2 ^82, 83^ predicted structure of MobA using anvi’o structure ^82–84^. The predicted catalytic domain of pBI143 MobA was structurally similar to MobM of the MobV-family (Protein Data Bank accession: 4LVI) encoded by plasmid pMV158 ^81^. We used the structurally similar catalytic domain in MobA to model the binding of the oriT of pBI143 to MobA. We found that there were only 21 SAAVs throughout MobA that were present in greater than 5% of the gut metagenomes (Fig. 3D). Interestingly, highly prevalent SAAVs occurred exclusively near the DNA binding site (L56, E49, and A64), suggesting that that the non-synonymous variants we observe in the context of MobA may be involved in altering the DNA binding specificity for the oriT sequence ^81^ demonstrating the coevolution of the oriT with the MobA protein between distinct pBI143 versions. Additionally, we find it likely that the cluster of high prevalence variation at residues V251, A246, V239, T238, I235, and L234 (Supplementary Fig. 4B) may be driven by interactions with different host conjugation machinery for plasmid transfer. The functional implications of prevalent SAAVs given the structural context of the MobA gene highlight the role of adaptive processes on the evolution of pBI143 versions.

Unlike the individual gut metagenomes, the pBI143 sequences did not occur in a monoclonal fashion in sewage metagenomes as expected (Supplementary Table 7). Sewage metagenomes had, on average, 35 SNVs with a departure from consensus value of lower than 0.9, revealing the polyclonal nature of pBI143 in sewage (Fig. 3C, Supplementary Table 7). Similar to the individual gut metagenomes, most SNVs in sewage metagenomes (78.8%) occurred at a nucleotide position that was variable across at least one pair of the three pBI143 versions (Fig. 3B, Supplementary Table 7), suggesting that the majority of the variability in sewage is from the mixing of different versions of pBI143. However, the number of novel SNVs was much higher in sewage: 61.8% of sewage samples had greater than three SNVs that did not match a variable position in one of the three reference plasmids (Fig. 3B). Given the marked increase in the number of novel SNVs in sewage, it is likely there are additional but relatively rare versions of pBI143 in the human gut.

Overall, these results indicate that pBI143 has a highly restricted mutational landscape in natural habitats, frequently occurs as a monoclonal element in individual gut metagenomes, and the non-synonymous variants of MobA in the environment may be responsible for altering its DNA binding.

### pBI143 is vertically transmitted, its variants are more specific to individuals than their host bacteria, and priority effects best explain its monoclonality in most individuals

The largely monoclonal nature of pBI143 presents an interesting ecological question: how do individuals acquire it, and what maintains its monoclonality? Multiple phenomena could explain the monoclonality of pBI143 in individual gut metagenomes, including (1) low frequency of exposure (i.e., most individuals are only ever exposed to one version), (2) bacterial host specificity (i.e., some plasmid versions replicate more effectively in certain bacterial hosts), or (3) priority effects (i.e., the first version of pBI143 establishes itself in the ecosystem and excludes others). The sheer prevalence and abundance of pBI143 across industrialized populations renders the ‘low frequency of exposure’ hypothesis an unlikely explanation. Yet the remaining two hypotheses warrant further investigation.

Bacterial host specificity is a plausible driver for the presence of a singular pBI143 version within an individual, given the interactions between plasmid replication genes and host replication machinery ^28, 85^. However, our analysis of 82 bacterial cultures isolated from 10 donors shows that the plasmid is more specific to individuals than it is to certain bacterial hosts (Fig. 4, Supplementary Table 9). Indeed, identical pBI143 sequences often occurred in multiple distinct taxa isolated from the same individual, in agreement with the monoclonality of pBI143 in gut metagenomes and its ability to transfer within Bacteroidales. If pBI143 monoclonality is not driven by rare exposure or host specificity, it could be driven by priority effects ^86^, where the initial pBI143 version somehow prevents other pBI143 versions from establishing in the same gut community.

**Fig. 4.**
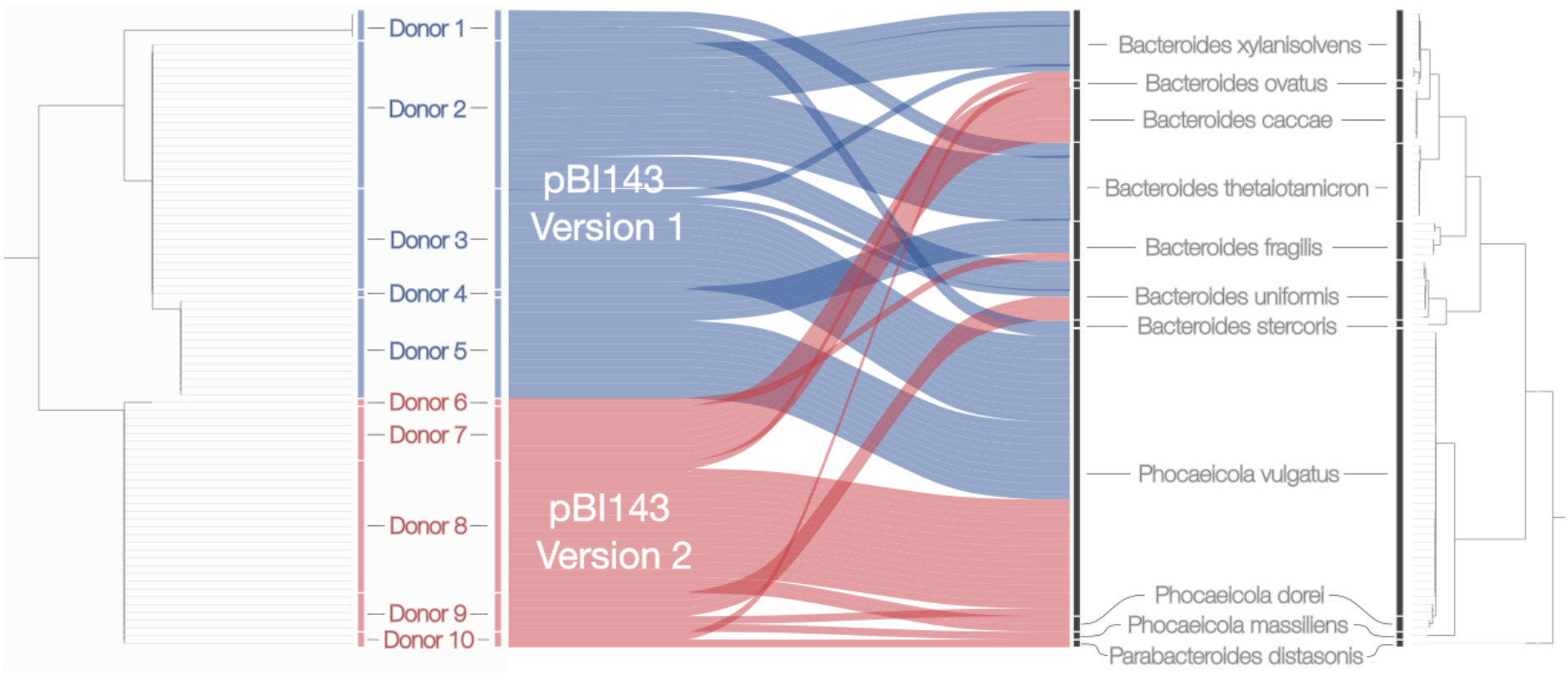
Phylogeny of pBI143 in human donors versus the phylogeny of bacterial isolates recovered from the same individuals. pBI143 (left) and bacterial host (right) genome phylogenies. The pBI143 phylogeny was constructed using the MobA and RepA genes; the bacterial phylogeny was constructed using 38 ribosomal proteins (see Methods). Blue alluvial plots are isolates with Version 1 pBI143 and red alluvial plots are isolates with Version 2 pBI143. No isolates had the rarer Version 3.

To examine if priority effects play a role in pBI143 monoclonality, we aimed to determine how pBI143 is acquired. Given that one established route of microbial acquisition is the vertical transmission of microbes from mother to infant ^87^, we used our ability to track pBI143 SNVs between environments to investigate if there is evidence for vertical transmission. We followed the inheritance of identical SNV patterns in pBI143 using 154 mother and infant gut metagenomes from four countries, Finland ^57^, Italy ^60^, Sweden ^68^, and the USA ^69^, where each study followed participants from birth to 3 to 12 months of age. We recruited reads from each metagenome to Version 1 pBI143 (Supplementary Table 1 and 3, Supplementary Fig. 5) and identified the location of each SNV in *mobA* (Supplementary Table 10). These data revealed a large number of cases where pBI143 had identical SNV patterns in mother-infant pairs (Fig. 5A, Supplementary Table 10). A network analysis of shared SNV positions across metagenomes appeared to cluster family members more closely, indicating mother-infant pairs had more SNVs in common than they had with unrelated individuals, which we could further confirm by quantifying the relative distance between each sample to others (Supplementary Fig. 6, Supplementary Table 10, Methods).

**Fig. 5.**
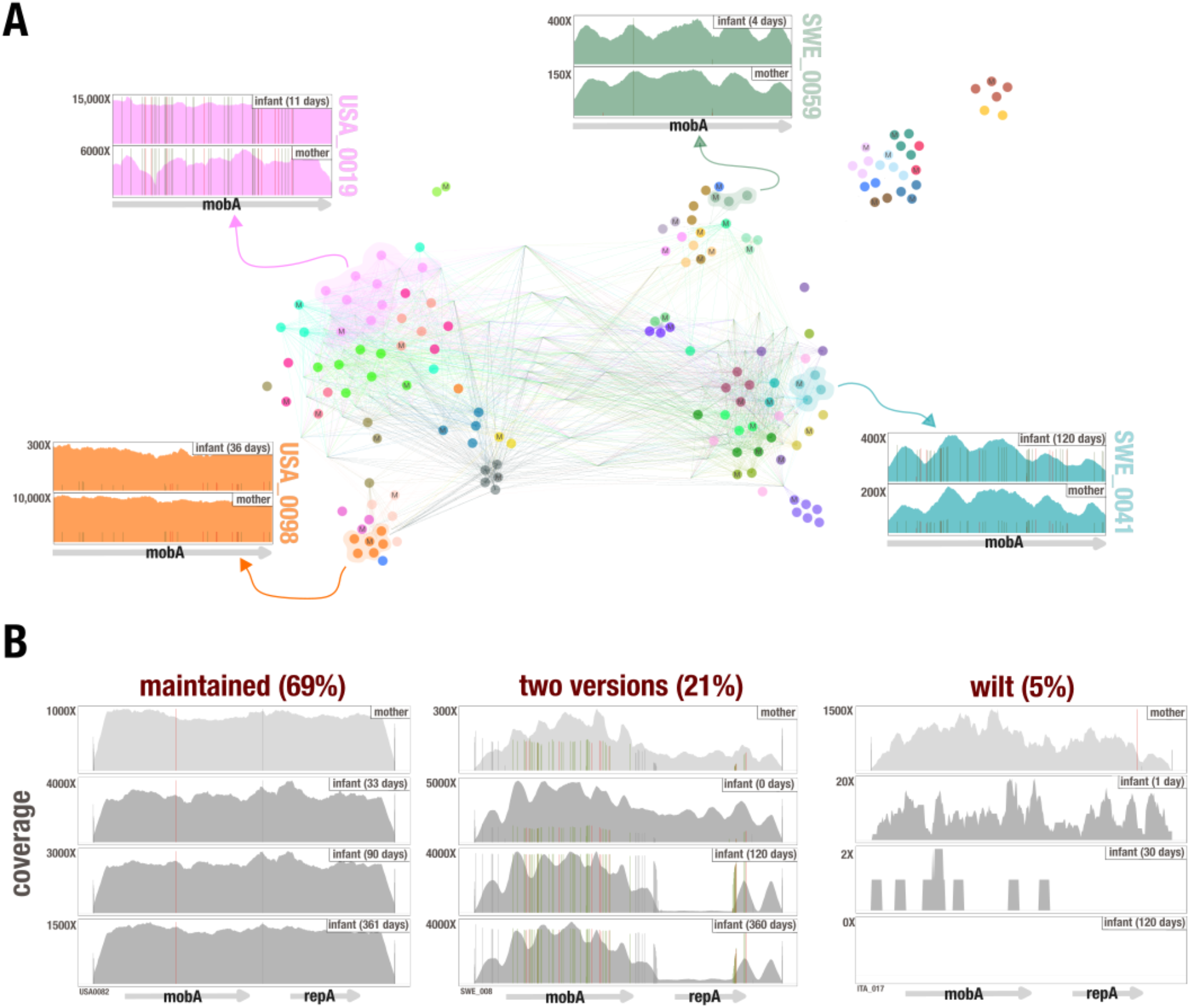
Transfer and maintenance of pBI143. **(A)** The network shows the degree of similarity between pBI143 SNVs across 154 mother and infant metagenomes from Finland, Italy, Sweden and the USA. Each node is an individual metagenome and nodes are colored based on family grouping. The surrounding coverage plots (colored) are visual representations of SNV patterns present in the indicated metagenomes. Nodes labeled with an “M” are mothers; nodes with no labels are infants. **(B)** Representative coverage plots showing different coverage patterns (maintained, two versions or wilt) observed in plasmids transferred from mothers to infants.

Establishing that pBI143 is often vertically transferred, we next examined the impact of priority effects on pBI143 maintenance over time. We assumed that if priority effects are driving persistence of a single version of pBI143, the first version that enters the infant gut environment should be maintained over time. Indeed, many phage populations are influenced by priority effects where the presence of one phage provides a competitive advantage to the host ^88^ or host immunity to infection with similar phages ^89–91^. In our data, we found no instances where pBI143 acquired from the mother was fully replaced in the infant during and up to the first year of life (Supplementary Fig. 5, Supplementary Table 10). While 69% of infants maintained the version received from the mother (Fig. 5B), we also observed other, less common genotypes. These less common cases included a ‘two versions’ scenario where the mother possessed two versions of pBI143, both of which were passed to the infant (21%), and a ‘wilt’ case, where the transferred pBI143 was neither replaced nor persisted until the end of sampling (7%) (Fig. 5B). Although these less prevalent phenotypes are not necessarily explained by priority effects, 69% maintenance of the initial version of pBI143 suggests that priority effects have an important role in the maintenance of pBI143 in the gut, despite many incoming populations colonizing the infant and likely carrying other pBI143 versions.

Overall, by tracking SNV patterns between environments we established that pBI143 is vertically transferred from mothers to infants and that priority effects likely play a role in maintaining the predominantly monoclonal populations of pBI143.

### pBI143 is a highly efficient parasitic plasmid

An intuitive interpretation of the surprising levels of prevalence and abundance of pBI143 across the human population, in addition to its limited variation maintained by strong evolutionary forces, is that it provides some benefit to the bacterial host. However, the two annotated genes in pBI143 appear to serve only the purpose of ensuring its own replication and transfer, contradicting this premise. The coverage of pBI143 and its *Bacteroides, Phocaeicola* and *Parabacteroides* hosts in gut metagenomes indeed show a significant positive correlation (R^2^: 0.5, p-value < 0.001) (Fig. 6A, Supplementary Table 11), however, these data are not suitable to distinguish whether pBI143 provides a benefit to the bacterial host fitness, or acts as a genetic hitchhiker.

**Fig. 6.**
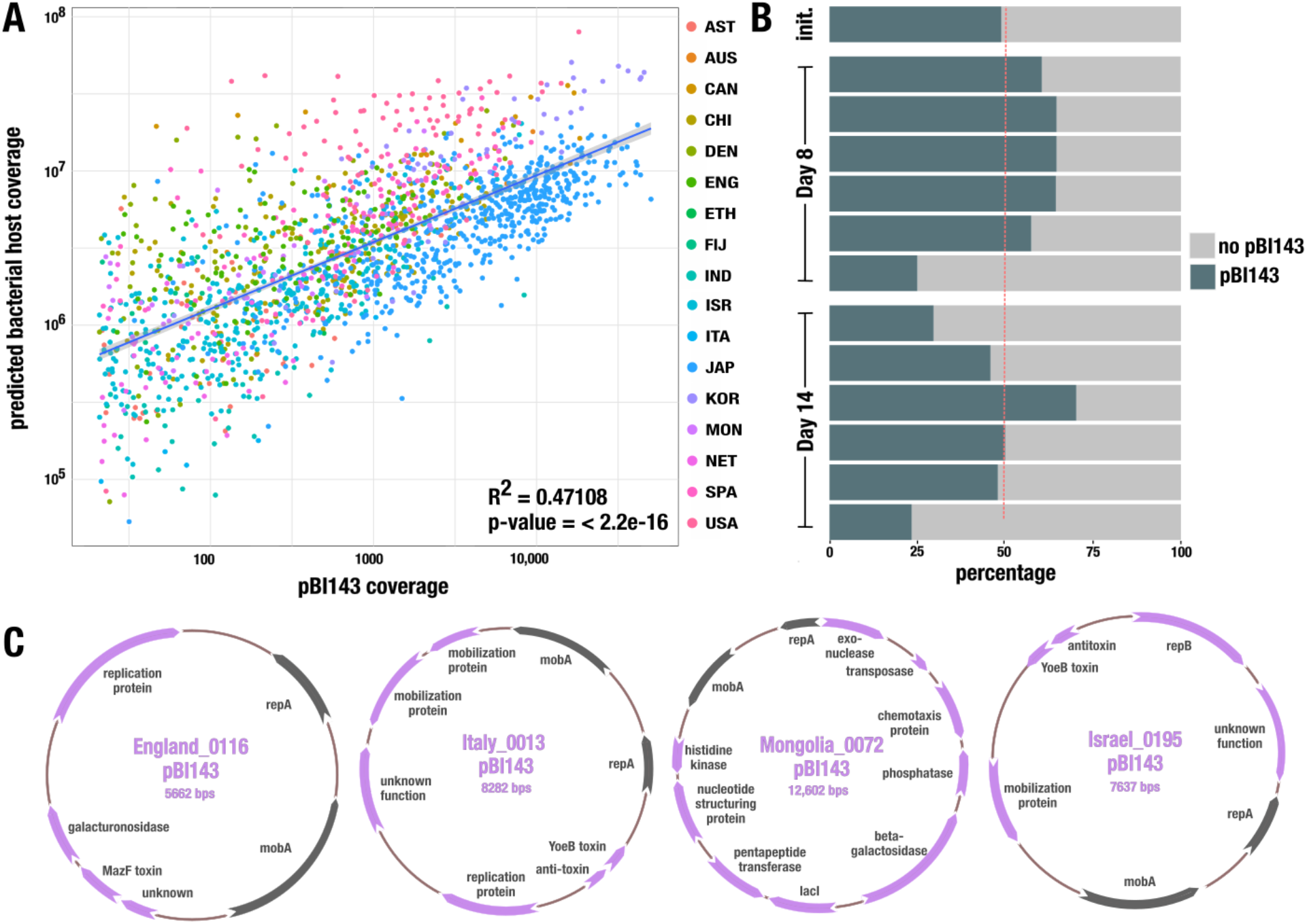
The relationship between pBI143 and its bacterial hosts. **(A)** The average coverage of pBI143 and the corresponding coverage of predicted host genomes (*Bacteroides, Parabacteroides and Phocaeicola*) in 4,516 metagenomes. **(B)** Competition experiments in gnotobiotic mice between *B. fragilis* with and without pBI143. The proportion of pBI143-carrying cells in 6 mice in the initial inoculum, at Day 8 and at Day 14 are shown. **(C)** Four examples of pBI143 assembled from metagenomes that carry additional cargo genes. Gray genes are the canonical *repA* and *mobA* genes of naive pBI143; lilac genes are additional cargo.

To experimentally investigate if pBI143 is advantageous or parasitic, we constructed isogenic pairs of *B. fragilis* 638R and *B. fragilis* 9343 with and without the native Version 1 sequence of pBI143 (Supplementary Methods). To determine if pBI143 is well-adapted to replication in a new *Bacteroides* host, we tested its maintenance in culture. After 7 days of passaging, pBI143 was still present in all colonies of *B. fragilis* 638R and *B. fragilis* 9343 (Supplementary Table 12). Next, we competed the *B. fragilis* 638R (with and without pBI143) of *B. fragilis* 638R in gnotobiotic mice for 2 weeks. At Day 8, 5/6 mice had more *B. fragilis* 638R with pBI143 than without; however this trend did not continue into Day 14, where 4/6 mice had fewer cells with pBI143 (Fig. 6B, Supplementary Table 12). While we can speculate that these populations may continue to fluctuate, the results at least suggest a negligible negative fitness impact of pBI143 on its bacterial host.

One potential benefit that pBI143 could provide to its host is to act as a natural shuttle vector by transiently acquiring additional genetic material and transferring it between cells in a community. In fact, in our survey of assembled gut metagenomes we observed a few cases that may support such a role for pBI143. In most individuals, we assembled pBI143 in its native form with 2 genes. However, there were 10 instances where the assembled pBI143 sequence from a given metagenome contained additional genes (Fig. 6C, Supplementary Table 2). Many of the additional genes have no predicted function, but other cargo include toxin-antitoxin genes conferring plasmid stability, as well as those that may confer beneficial functions to the bacterial host, such as galacturonosidase, pentapeptide transferase, phosphatase, and histidine kinase genes. These occasional larger versions of pBI143 share a common backbone of *repA* and *mobA* and thus form a “plasmid system” ^40^, a common plasmid evolutionary pattern suggesting the possibility that pBI143 may dynamically acquire different genes in different environments.

Overall, it does not appear that the native sequence of pBI143 provides a clear benefit to its host cells, however it does appear to positively correlate with these hosts in metagenomic data, and is maintained in the absence of selection in new hosts *in vitro*.

### pBI143 responds to oxidative stress *in vitro*, and its copy number is significantly higher in metagenomes from individuals who are diagnosed with IBD

Mobile genetic elements rely on their hosts for replication machinery, but many have developed mechanisms to increase their rates of replication and transfer during stressful conditions to increase the likelihood of their survival if the host cell dies ^92–95^. To investigate whether the copy number of pBI143 changes as a function of stress, we first conducted an experiment with *B. fragilis* isolates that naturally carry pBI143.

Given that oxygen exposure upregulates oxidative stress response pathways in the anaerobic *B. fragilis* ^96^, we exposed two different *B. fragilis* cultures, *B. fragilis* R16 (which was isolated from a healthy individual) and *B. fragilis* 214 (which was isolated from a pouchitis patient ^97^) to 21% oxygen for increasing periods of time (Fig. 7A, Supplementary Fig. 7, Supplementary Table 13). To calculate the copy number of pBI143 in culture, we quantified the ratio between the total number of plasmids and the total number of cells in culture using a qPCR with primers targeting pBI143 and a *B. fragilis*-specific gene we identified through pangenomics (Methods). As the length of oxygen exposure increased, the copy number of pBI143 per cell also increased. Notably, the copy number was quickly reduced to control levels once the cultures were returned to anaerobic conditions, indicating that copy number fluctuation is a rapid and transient process that is dependent on host stress.

**Fig. 7.**
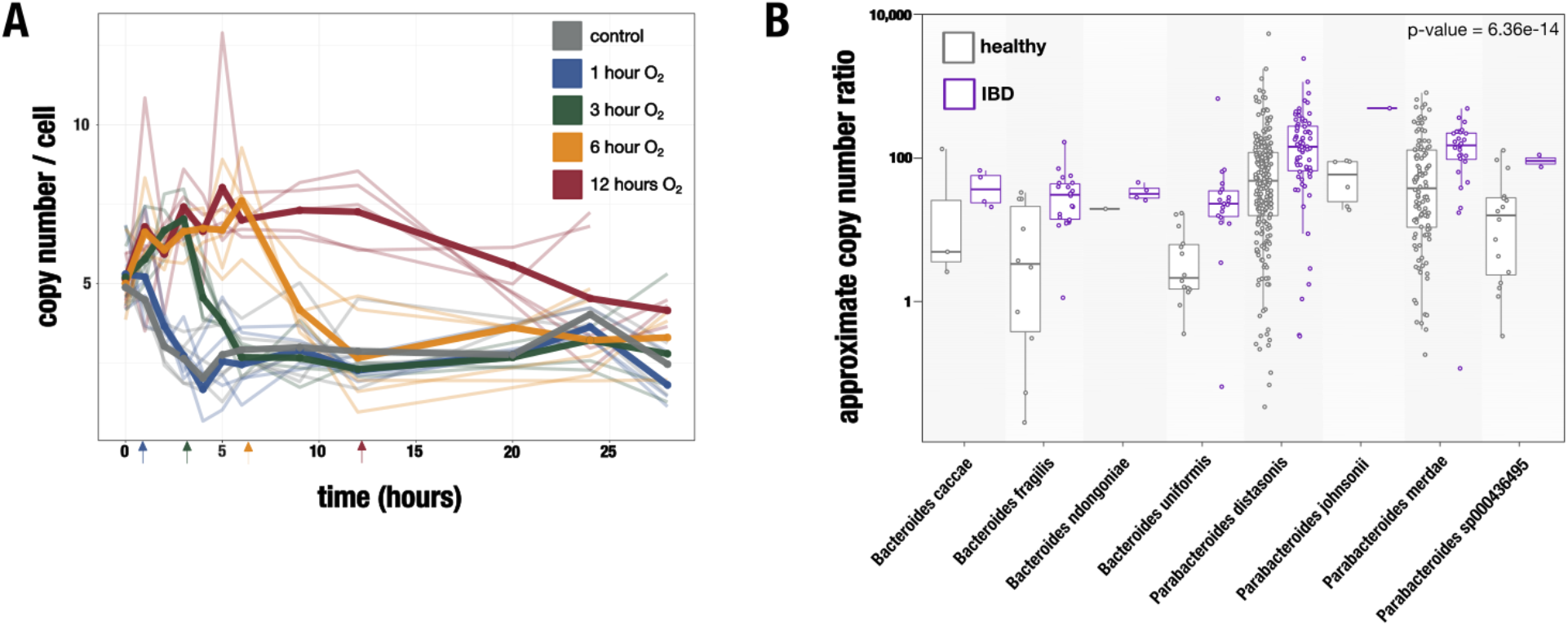
pBI143 copy number increases in stressful environments. **(A)** Copy number of pBI143 in *B. fragilis* 214 cultures with increasing exposure to oxygen. Arrows indicate the time point at which the culture was returned to the anaerobic chamber. The control cultures (gray) were never exposed to oxygen. Opaque lines are the mean of 5 replicates (translucent lines). **(B)** Host-specific approximate copy number ratio (ACNR) of pBI143 in healthy individuals (gray) versus those with IBD (purple).

Oxidative stress is also a signature characteristic of inflammatory bowel disease (IBD), a group of intestinal disorders that cause inflammation of the gastrointestinal tract ^98^. The dysregulation of the immune system during IBD typically leads to high levels of oxidative stress in the gut environment ^99^. We thus hypothesized that, if oxidative stress is among the factors that drive the increased copy number of pBI143 in culture, one should expect a higher copy number of pBI143 in metagenomes from IBD patients compared to healthy controls.

To analyze the copy number of pBI143 in a given metagenome, we calculated the ratio of metagenomic read coverage between pBI143 and its bacterial host in metagenomes where pBI143 could confidently be assigned to a single host. With these considerations, we developed an approach to calculate an ‘approximate copy number ratio’ (ACNR) for pBI143 and its unambiguous bacterial host in a given metagenome using bacterial single-copy core genes (see Methods). We calculated the ACNR of pBI143 in 3,070 healthy and 1,350 IBD gut metagenomes (Supplementary Table 1, Supplementary Fig. 1 and 8). Our analyses showed that the geometric mean of the ACNR for pBI143 and its host was 3.72 times larger (robust-Wald 95% CI: 2.66x - 5.20x, p-value < 10^-13^) in IBD compared to healthy metagenomes, indicating that the pBI143 ACNR was significantly higher in individuals with IBD compared to those who were healthy (Fig. 7B, Supplementary Table 14).

The copy number ratio of pBI143 to its *B. fragilis* host in culture calculated with qPCR primers was much lower (∼5X on average) compared to its approximate copy number ratio in healthy metagenomes (∼120X on average). Multiple factors can explain this difference, including biases associated with sequencing steps or the calculation of the coverage, or that the conditions naturally occuring communities experience vastly differ than those conditions encountered in culture media, even in the presence of oxygen. Nevertheless, the marked increase of the relative coverages of pBI143 and its host in IBD metagenomes suggest the potential utility of this cryptic plasmid for unbiased measurements of stress. Overall, these results show that both in metagenomes and experimental conditions, an increased copy number of pBI143 is a consistent phenotype in the presence of host stress.

## DISCUSSION

Our work shed lights on a mysterious corner of life in the human gut. Even though pBI143 is found in greater than 90% of all individuals in some countries, the prevalence of this cryptic plasmid has gone unnoticed for almost four decades since its discovery by Smith, Rollins, and Parker ^41^. The remarkable ecology, evolution, and potential practical applications of pBI143 that we characterized here through ‘omics analyses as well as *in vitro* and *in vivo* laboratory experiments offer a glimpse of the world of understudied cryptic plasmids in the human gut, and elsewhere.

The application of population genetics principles to pBI143 through the recovery of single-nucleotide variants (SNVs) and single-amino acid variants (SAAVs) from gut metagenomes reveals not only the strong forces of purifying selection on the evolution of its sequence, but also hints the presence of adaptive processes at localized amino acid positions that are variable in the critical parts of the DNA-interacting residues of the catalytic domain of its mobilization protein. The presence of pBI143 does not appear to systematically impact bacterial host fitness *in vivo*, which makes this cryptic plasmid seem a mundane parasite, somewhat contradicting the strict evolutionary pressures that maintain its environmental sequence variants.

That said, our observations from naturally occurring gut environments include cases where pBI143 carries additional genes, likely acting as a natural shuttle vector. Although traditionally mobile genetic elements are classified as mutualistic or parasitic with respect to the bacterial host, the fluidity of pBI143 to fluctuate between the cryptic 2-gene state and the larger 3 or more gene state with potentially beneficial functions, suggests that the boundaries between parasitism and mutualism for pBI143 are not clear cut. Instead, pBI143 may act as a ‘discretionary parasite’, where it has a cryptic form for the majority of its existence in which it could be best described as a parasite, while occasionally being found with additional functions that may be beneficial to its host as a function of environmental pressures. Testing this hypothesis with future experimentation, and if true, investigating to what extent discretionary parasitism applies to cryptic plasmids, may lead to a deeper understanding of the role cryptic plasmids play in microbial fitness to changing environmental conditions.

Our findings show that it has important potential practical applications beyond molecular biology. The first and most straightforward of these applications relies on the prevalence and human specificity of pBI143 to more sensitively detect human fecal contamination in water samples. Human fecal pollution is a global public health problem, and accurate and sensitive indicators of human fecal pollution are essential to identify and remediate contamination sources and to protect public health ^100^. While culture assays for *E. coli* or enterococci have historically been used to detect human fecal contamination in environmental samples, the common occurrence of these organisms in many different mammalian guts and the poor sensitivity of such assays motivated researchers in the past two decades to utilize PCR amplification of 16S rRNA genes, specifically those from human-specific *Bacteroides* and *Lachnospiraceae* populations, to detect human-specific fecal contamination with minimal cross-reactivity with animal feces ^79, 80^. Our benchmarking of pBI143 with qPCR revealed that pBI143 is an extremely sensitive and specific marker of human fecal contamination that typically occurs in human fecal samples and sewage in numbers that are several-fold higher than the state-of-the-art markers, which enabled the quantification of fecal contamination in samples where it had previously gone undetected. Another practical application of pBI143 takes advantage of its natural shuttle vector capabilities to incorporate additional genetic material into its backbone. Our demonstration that pBI143 (1) replicates in many abundant gut microbes, (2) can be stably introduced to new hosts, and (3) naturally acquires genetic material makes this cryptic plasmid an ideal natural payload delivery system for future therapeutics targeting the human gut microbiome. Indeed, our observations of pBI143 with cargo genes in metagenomes indicates that this likely happens in nature. Yet another practical implication of pBI143 is its utility to measure the level of stress in the human gut. Surveying thousands of samples from individuals who are healthy or diagnosed with IBD, our results show that across all bacterial hosts, the approximate copy number of pBI143 increases in individuals with IBD.

From a more philosophical point of view, the prevalence and high conservancy of pBI143 across globally distributed human populations questions the traditional definition of the ‘core’ microbiome ^101^. In its aim to define a core microbiome, the field of microbial ecology has primarily focused on bacteria, although sometimes including prevalent archaea or fungi ^102–105^. However, our results indicate that there are mobile genetic elements that fit the standard criteria of prevalence to be defined as core. Broadening the definition of a core microbiome beyond microbial taxa may enable the recognition of other mobile genetic elements (eg. plasmids, phages, transposons) that are prevalent across human populations and fill critical gaps in our understanding of gut microbial ecology.

## Materials and Methods

### Genomes and metagenomes

We acquired the original pBI143 genome from the National Center for Biotechnological Information (GenBank: U30316.1). We manually assembled the three reference versions of pBI143 (Version 1, 2 and 3) from metagenomes samples USA0006, CHI0054 and ISR0084. We acquired 717 human gut isolate genomes from the Duchossois Family Institute collection (Supplementary Table 4). We downloaded 4,516 healthy human adult gut metagenomes from the National Center for Biotechnology Information (NCBI) from (Australia (Accession ID: PRJEB6092), Austria ^48^, Bangladesh ^49^, Canada ^50^, China ^51, 52^, Denmark ^53^, England ^54^, Ethiopia ^55^, Fiji ^56^, Finland ^57^, India ^58^, Israel ^59^, Italy ^60, 61^, Japan ^62^, Korea ^63^, Madagascar ^55^, Mongolia ^55, 64^, Netherlands ^65^, Peru ^66^, Spain ^67^, Sweden ^68^, Tanzania ^61^, and the USA ^66, 69, 70^) (Supplementary Table 1). We acquired 1,096 gut metagenomes from infant-mother pairs from Italy, Finland, Sweden and the USA from NCBI (Supplementary Table 1). We downloaded 935 metagenomes from non-human gut environments (marine ecosystems, pet dog guts, monkey guts, sewage, human oral cavity, and human skin) (Supplementary Table 1).

### Metagenomic assembly, read recruitment, and the recovery of coverage and detection statistics

Unless otherwise specified, we performed all metagenomic analyses throughout the manuscript within the open-source anvi’o v7 software ecosystem (https://anvio.org) ^106^. We automated assembly and read recruitment steps using the anvi’o metagenomics workflow ^107^ which used snakemake v5.10 ^108^. To quality-filter genomic and metagenomic raw paired-end reads we used illumina-utils v1.4.4 ^109^ program ‘iu-filter-quality-minoch’ with default parameters, and IDBA_UD v1.1.2 with the flag ‘--min_contig 1000’to assemble the metagenomes ^110^. We used Bowtie2 v2.4 ^111^ to recruit reads from the metagenomes to reference sequences and samtools v1.9 ^112^ to convert resulting SAM files into sorted and indexed BAM files. We generated anvi’o contigs databases (https://anvio.org/m/contigs-db) using the command ‘anvi-gen-contigs-databasè, during which Prodigal v2.6.3 ^113^ identifies open reading frames. We created anvi’o profile databases of the mapping results for each metagenome using ‘anvi-profilè, which stores coverage and detection statistics, and ‘anvi-merg’ to combine all profiles together. To recover coverage and detection statistics for a given merged profile database, we used the program ‘anvi-summariz’ with ‘--init-gene-coverages’flag.

### Criteria for detection of pBI143 and crAssphage in metagenomes

Using mean coverage to assess the occurrence of a given sequence in a given sample based on metagenomic read recruitment can yield misleading insights due to non-specific read recruitment (i.e., recruitment of reads from metagenomes to a reference sequence from non-target populations). Thus, we relied upon the detection statistic reported by anvi’o, which is a measure of the proportion of the nucleotides in a given sequence that are covered by at least one short read. We considered pBI143 was present in a metagenome only if its detection value was 0.5 or above. Values of detection in metagenomic read recruitment results often follow a bimodal distribution for populations that are present and absent (see Supplementary Fig. 2 in ref. ^114^). Thus, 0.5 is a conservative cutoff to minimize a false-positive signal to assume presence.

### Distinguishing the presence of distinct pBI143 versions in a genome or metagenome

We used the results of individual read recruitments to each known version of pBI143 to measure the coverage of each gene in pBI143 in samples that had a detection of greater than 0.9 and compared the ratio of the coverage of each gene. The pBI143 version where the genes have the most even coverage ratio was considered the predominant version in that genome or metagenome.

### Addition of tetQ to pIB143

To study transfer of pBI143 from *Phocaeicola vulgatus* MSK 17.67 to other Bacteroidales species, we added *tetQ* to pBI143. We PCR amplified tetQ from *Bacteroides caccae* CL03T12C61 and inserted it at the site shown in Supplementary Fig. 2 (all primers are listed in Supplementary Table 15). We PCR amplified the DNA regions flanking each side of this insertion site and the three PCR products were cloned into BamHI-digested pLGB13 ^115^. We conjugally transferred this plasmid into *Phocaeicola vulgatus* MSK 17.67 and selected cointegrates on gentamycin 200 µg/ml and erythromycin 10 µg/ml. We passaged the cointegrate in non-selective medium and selected the resolvents by plating on anhydrotetracycline (75 ng/ml). We confirmed pIB143 contained *tetQ* by WGS the strain at the DFI Microbiome Metagenomics Facility.

### Transfer assays

The recipient strains that received pBI143-*tetQ* were *Parabacteroides johnsonii* CL02T12C29 and *Bacteroides ovatus* D2, both erythromycin resistant and tetracycline sensitive. We grew the donor strain *Phocaeicola vulgatus* MSK 17.67 pBI143-*tetQ* and recipient strains to an OD600 of ∼ 0.7 and mixed them at a 10:1 ratio (v:v) donor to recipient, and spotted 10 µl onto BHIS plates and grew them anaerobically for 20 h. We resuspended the co-culture spot in 1 mL basal media and cultured 10-fold serial dilutions on plates with erythromycin (to calculate number of recipients) or erythromycin and tetracycline (4.5 µg/ml) (to select for transconjugants). We performed multiplex PCR as described ^116, 117^ to confirm that TetR ErmR colonies were the recipient strain containing pBI143-tetQ (Supplementary Fig. 2).

### Calculations of purifying selection and characterization of single nucleotide variants across metagenomes

We calculated dN/dS ratios as described previously ^84^; details of which can also be found at https://merenlab.org/data/anvio-structure/chapter-IV/#calculating-dndstextgene-for-1-gene. To determine the mutational landscape of pBI143 across metagenomes, we first identified all variable positions present in the reference pBI143 sequences. We used the program ‘anvi-script-gen-short-reads’to generate artificial short reads from the version 2 and version 3 pBI143 sequences and recruited these reads to the version 1 pBI143 sequence to generate data similar to the read recruitment from metagenomes. Then, we took read recruitment data from the global human gut metagenomes and sewage metagenomes mapped to version 1 pBI143. We ran ‘anvi-gen-variability-profil’ on the artificial read recruitment profile databases as well as on all profile databases from metagenomes with greater than 10X Q2Q3 coverage to identify all SNV positions. We compared the SNV positions in each gut or sewage metagenome to those present in our reference sequences and calculated the number of SNVs in each metagenome that did and did not match SNVs in the references. To calculate the number of non-consensus SNVs in a metagenome, we again ran the command ‘anvi-gen-profile-databas’ on the same metagenomes, this time with the flags ‘--gene-caller- ids 0’, ‘--min-departure-from-consensus 0.1’, ‘--include-contig-names’and ‘--quince-mod’, which produces a file that describes the variation in every single position across the reference and calculates the departure from consensus for each SNV with a departure from consensus greater than 0.1.

### pBI143 structural and polymorphism analysis

To explore the impact of SAAVs on the protein structure of pBI143 MobA, we *de novo* predicted the monomer and dimer structures using AlphaFold 2 (AF) in ColabFold with default settings ^83^. AlphaFold 2 confidently predicted the structure of the catalytic domain but had low pLDDT scores for the coil domains and the dimer interactions. However, we explored variants across the whole dimer complex. Next, we integrated the pBI143 MobA AF structure into anvi’o structure by running ‘anvi-gen-structure-database’ ^82^. After that, we summarized SNV data as SAAVs from the metagenomic read recruitment data using ‘anvi-gen-variability-profile --engine AA’ to create a variability profile (https://anvio.org/m/variability-profile). Subsequently, we superimposed the SAAV data variability profile on the structure with ‘anvi-display-structure’ which filtered for variants that had at least 0.05 departure from consensus (reducing our metagenomic samples size from 2221 to 1706). Finally, we analyzed SAAVs that were prevalent in at least 5% of remaining samples. This left us with 21 SAAVs to analyze on the monomer. Next, we explored the relationship between SAAVs, relative solvent accessibility (RSA), and ligand binding residues in pBI143 MobA. To do this, we identified the homologous structure PDB 4LVI (MobM) by searching the high pLDDT pBI143 AF domain against the structure database PDB100 2201222 using Foldseek (https://search.foldseek.com/search). We next structurally aligned the pBI143 MobA AF structure to PDB 4LVI (MobM) ^118^ using PyMol ^119^. We chose the MobM structure 4LVI rather than a MobA because it had more structural and sequence homology to the pBI143 MobA catalytic domain AF structure than any PDB MobA structures. Additionally, we leveraged residue conservation values from the pre-calculated <u>4LVI ConSurf analysis</u> to further explore ligand binding residues ^120, 121^.

### Phylogenetic tree construction

To construct the pBI143 phylogeny, we identified pBI143 contigs from the isolate genome assemblies (Supplementary Table 4) using BLAST ^122^. We ran ‘anvi-gen-contigs-database’ on each pBI143 contig followed by ‘anvi-export-gene-calls’ with the flag ‘--gene-caller prodigal’ and concatenated the resulting amino acid sequences. For the bacterial host phylogeny, we ran ‘anvi-gen-contigs-database’ on each assembled genome. Then, we extracted ribosomal genes (see Supplementary Methods for details), aligned them with MUSCLE v3.8.1551 ^123^, trimmed the alignments with trimAl ^124^ using the flag ‘-gt 0.5’, and computed the phylogeny with IQ-TREE 2.2.0-beta using the flags ‘-m MFP’ and ‘-bb 1000’ ^125^. We visualized the trees with ‘anvi-interactive’ in ‘--manual-mode’, and used the metadata provided by the Duchossois Family Institute to label the isolates to their corresponding donors. We used the ‘geom_alluvium’function in ggplot2 to make the alluvial plots..

### Construction and analysis of the network that describes shared single-nucleotide variants across mothers and infants

To investigate whether single-nucleotide variants suggest a vertical transmission of pBI143, we used metagenomic read recruitment results from four independent study that generated metagenomic sequencing of fecal samples collected from mothers and their infants in Finland ^57^, Italy ^60^, Sweden ^68^, and the USA ^69^, against the pBI143 Version 1 reference sequence. The URL https://merenlab.org/data/pBI143 serves a fully reproducible workflow of this analysis. The primary input for this investigation was the anvi’o variability data, which is calculated by the anvi’o program ‘anvi- profil’, and reported by the anvi’o program ‘anvi-gen-variability-profil’ (with the flag ‘--engine NT’). The program ‘anvi-gen-variability-profil’ (https://anvio.org/m/anvi-gen-variability-profile) offers a comprehensive description of the single-nucleotide variants in metagenomes for downstream analyses. Since the *mobA* gene was conserved enough to represent all three versions of pBI143, for downstream analyses we limited the context to study variants to the mobA gene. The total number of samples in the entire dataset with at least one variable nucleotide position was 309, which represented a total of 102 families (Sweden: 52, USA: 24, Finland: 14, Italy: 11). We removed any sample that did not belong to a minimal complete family (i.e., at least one sample for the mother, and at least one sample of her infant), which reduced the number of families in which both members are represented to 57 families (Sweden: 36, USA: 16, Finland: 3, Italy: 2). We further removed families if the coverage of the *mobA* gene was not 50X or more in at least one mother and one infant sample in the family, which reduced the number of families with both members represented and with a reliable coverage of mobA to 49 families (Sweden: 33, USA: 13, Finland: 2, Italy: 1), and from a given family, we only used the samples that had at least 50X for downstream analyses. We subsampled the variability data in R to only include the variable nucleotide position data for the final list of samples. We then used the list of single-nucleotide variants reported in this file to generate a network description of these data using the program ‘anvi-gen-variability-network’, which reports an ’edge’ between any sample pairs that share a SNV with the same competing nucleotides. We then used Gephi ^126^, an open-source network visualization program, with the ForceAtlas2 algorithm ^127^ to visualize the network. To quantify the extent of similarity between family members based on single-nucleotide patterns in the data, we generated a distance matrix from the same dataset using the ‘pdist’function in Python’s standard library with ‘cosin’ distances. We calculated the average distance of each sample to all other samples in its familial group (‘within distanc’), as well as the average distance from each sample to all other samples not present in their familial group (‘between distanc’). We subtracted the within distance from the between distance to get the ‘subtracted distancè.

### Metagenomic taxonomy estimation

We used Kraken 2.0.8-beta with the flags ‘--output’, ‘--report’, ‘-- use-mpa-styl’, ‘--quick’, ‘--use-names’, ‘--paired’and ‘--classified-out’to estimate taxonomic composition of each metagenome ^128^. For the genus-level taxonomic data, we filtered for metagenomes where the total number of reads recruited to a *Bacteroides*, *Parabacteroides* or *Phocaeicola* genome was >1000 and the mean coverage of pBI143 was >20X. For the species-level taxonomic data, we used a cutoff of >0.1% percent of reads recruited to designate presence or absence of *B. fragilis* and >0.0001% for pBI143 based on the sizes of the genomes respectively (the *B. fragilis* genome is 3 orders of magnitude larger than pBI143).

### Isogenic strain construction

We constructed the plasmid vector pEF108 (as shown in Supp Fig. pEF108_plasmid_map) by PCR amplifying the desired sections with primers vec_108F, vec_108R, frag1_108F, frag1_108R, frag2_108R and frag2_108R (Supplementary Table 15) from existing plasmids. We assembled the three fragments via Gibson assembly using standard conditions described for NEB Gibson assembly mastermix. We selected for transconjugants on LB-carbenicillin (100ug/mL), then conjugated pEF108 into B. fragilis 638R and selected on BHIS + erythromycin 25ug/mL. Then, we counter-selected for recombination events in pEF108 to remove the markers and leave naive pBI143 by growing cells on Bacteroides minimal media plates (BMM) with 10mM p-chlorophenylalanine. We screened pBI143 positive, pheS-negative colonies via PCR and confirmed them by WGS. See Supplementary Methods for details.

### *In vitro* competition experiments

We grew each strain described above in BHIS to OD 0.6-0.8. We combined equal volumes of cells and plated these cells on BHIS plates. We added 50µL of the combined strains to 5mL BHIS and grew these cultures to OD 0.6-0.8, then plated again on BHIS plates. We replica plated all colonies from BHIS to BHIS supplemented with cefoxitin (15ug/mL) or erythromycin (10 ug/mL) and counted the resulting colonies to determine the starting and final ratios of each strain.

### Mouse competitive colonization assays

All animal experimentation was approved by the Institutional Animal Care and Use Committee at the University of Chicago. We gavaged three male and three female 10-15 week old germ-free C57BL/6J mice with a 1:1 inoculum of *B. fragilis* 638R:*B. fragilis* 638R pBI143. Males and females were housed separately in isocages and remained gnotobiotic for the duration of the experiment. We collected fecal pellets after eight and 14 days, diluted and plated on BHIS plates. We performed PCR on 48 colonies per mouse using a mixture of four primers (Supplementary Table 15), one set that amplifies a 1248-bp region of the 638R chromosome and a second set that amplifies a 662-bp segment of pBI143. PCR amplicons from all colonies included the 1248-bp region of the 638R chromosome and a subset also contained the amplicon for pBI143, allowing calculation of the ratio over time. The exact starting ratio for gavage was also calculated using this same PCR.

### Approximate copy number ratio calculation in metagenomes

The first challenge to use metagenomic coverage values to study pBI143 copy number trends in human gut metagenomes is the unambiguous identification of gut metagenomes that appear to have a single possible pBI143 bacterial host beyond reasonable doubt. To establish insights into the taxonomic make up of the gut metagenomes we previously assembled, we first ran the program ‘anvi-estimate-scg-taxonomy’(https://anvio.org/m/anvi-estimate-scg-taxonomy) with the flags ‘--metagenome-mod’ (to profile every single single-copy core gene (SCG) independently) and ‘--compute-scg-coverages’(to compute coverages of each SCG from the read recruitment results). We also used the flag ‘--scg-name-for-metagenome-mod’ to limit the search space for a single ribosomal protein. We used the following list of ribosomal proteins for this step as they are included among the SCGs anvi’o assigns taxonomy using GTDB, and we merged resulting output files: Ribosomal_S2, Ribosomal_S3_C, Ribosomal_S6, Ribosomal_S7, Ribosomal_S8, Ribosomal_S9, Ribosomal_S11, Ribosomal_S20p, Ribosomal_L1, Ribosomal_L2, Ribosomal_L3, Ribosomal_L4, Ribosomal_L6, Ribosomal_L9_C, Ribosomal_L13, Ribosomal_L16, Ribosomal_L17, Ribosomal_L20, Ribosomal_L21p, Ribosomal_L22,ribosomal_L24, and Ribosomal_L27A. For our downstream analyses that relied upon the merged SCG taxonomy and coverage output reported by anvi’o, we considered *Bacteroides, Parabacteroides* and *Phocaeicola* as the genera for candidate pBI143 host ‘species’, and only considered metagenomes in which a single species from these genera was present. Our determination of whether or not a single species of these genera was present in a given metagenome relied on the coverage of species-specific single-copy core genes (SCGs), where the taxonomic assignment to a given SCG resolved all the way down to the level of species unambiguously. We excluded any metagenome from further consideration if three or more candidate host species had positive coverage in any SCG in a metagenome. Due to highly conserved nature of ribosomal proteins and bioinformatics artifacts, it is possible that even when a single species is present in a metagenome, one of its ribosomal proteins may match to a different species in the same genus given the limited representation of genomes in public databases compared to the diversity of environmental populations. So, to minimize the removal of metagenomes from our analysis, we took extra caution with metagenomes before discarding them if only two candidate host species had positive coverage in any SCG. We kept such a metagenome in our downstream analyses only if one species was detected with only a single SCG, and the other one was detected by at least 8. In this case we assumed the large representation of one species (with 8 or more ribosomal genes) suggests the presence of this organism in this habitat confidently, and assumed the single hit to another species within the same genus was likely due to bioinformatics artifacts. It is the most unambiguous case if only a single candidate host species was detected in a given metagenome, but we still removed a given metagenome from further consideration if that single species had 3 or fewer SCGs in the metagenome. These criteria deemed 584 of 2580 metagenomes to have an unambiguous pBI143 host that resolved to 21 distinct species names. We further removed from our modeling the metagenomes where the candidate host species did not occur in any other metagenome, which removed 5 of these candidate host species from further consideration. Finally, we further removed any metagenome in which the pBI143 coverage was less than 5X. Our final dataset to calculate the “approximate copy number ratio” (ACNR) of pBI143 in metagenomes through coverage ratios contained 579 metagenomes with one of 16 unambiguous pBI143 hosts. We calculated the ACNR by dividing the observed coverage of pBI143 by the empirical mean coverage of the host by averaging the coverage of all host SCGs found in the metagenome. To estimate the multiplicative difference in the geometric mean ACNR, we fit a linear model for the expected value of the logarithm of the ACRN, with disease status and bacterial host as predictors using rigr to construct the interval and estimate ^129^ .

### Oxidative stress experiments

We grew *B. fragilis* 214 in 5 mL BHIS for 15 hours in an anaerobic chamber. We inoculated 750 µL of this culture into 30mL BHIS in quintuplicate, and grew them for 3 hours. We divided the 30 mL into a further 5 culture flasks of 5 mL BHIS, and exposed each to oxygen with constant shaking for the appropriate time before returning the flask to the anaerobic chamber. At each time point, we took an aliquot of culture to determine the copy number of pBI143 in that sample. We extracted DNA from the cultures using a Thermal NaOH preparation ^130^ to prepare them for qPCR. Copy number calculated can be found in Supplementary Table 13.

### Estimating the pBI143 Plasmid Copy Number by Real-time qPCR

To evaluate plasmid copy number (CN), we developed a real-time TaqMan probe multiplex PCR assay to amplify both pBI143 and a single-copy *B. fragilis*-specific genomic reference gene (referred to as hsp [heat shock protein]) in the same reaction (see Supplementary Information for details). We confirmed the primer and probe specificity to *B. fragilis* with BLAST searches against the NCBI and Ensembl databases, and experimental validation on 45 common gut isolates. For absolute quantification, we constructed a standard curve for each gene of interest by plotting the mean quantification cycle (Cq) values against log[quantity] of a dilution series of known gene of interest amount (range: 3×10^0 to 3×10^6 copies/reaction). We calculated the CN of pBI143 per genome equivalent (hsp), by dividing the absolute quantity of plasmid target by the absolute quantity of chromosomal target in the sample using the standard-curve (SC) method of absolute quantification ^131^. Standard curves were generated with every qPCR run for analysis and to confirm PCR efficiency. Additional details for qPCR, including standard curves and controls, can be found in Supplemental Information. Supplementary Table 5 and Supplementary Table 15 report the relevant data and all primers, respectively.

### qPCR analysis of animal, untreated sewage and water samples

Samples were tested with the pBI143 assay and two established assays for human fecal markers that included HF183 and Lachno3 ^132^. For screening of animal samples to assess the presence of this plasmid in non-human gut microbiomes, archived DNA from a previous study ^133^ was analyzed and included 14 different animals encompassing 81 individual fecal samples. For assessment of fecal contamination of surface waters, archived DNA from 40 samples of river water ^134–136^ and freshwater beaches ^137^ were analyzed. These water samples were chosen from these previous studies that represented a range of contamination based on HF183 and Lachno3 levels. A total of 20 archived untreated sewage samples as reported in Olds et al. ^132^ were also analyzed for comparison. Since we were using archived samples from previous studies, we retested all the samples for the two human markers to account for any degradation. Additional details for qPCR, including standard curves and controls, can be found in the Supplemental Information.

### Visualizations

We used ggplot2 ^138^ to generate all box and scatter plots. We generated coverage plots using anvi’o, with the program ‘anvi-script-visualize-split-coverages’. We finalized the figures for publication using Inkscape, an open-source vector graphics editor (available from http://inkscape.org/).

## Data availability

All genomes and metagenomes are available via the NCBI Sequence Read Archive, and the accession numbers for metagenomes and genomes are reported in Supplementary Table 1 and Supplementary Table 4, respectively. The data object identifier (DOI) <u>10.6084/m9.figshare.22336666</u> gives access to Supplementary Table and Supplementary Information files. Additional DOIs for anvi’o data products that describe metagenomic read recruitment results as well as sequences for pBI143 versions and bioinformatics workflows are accessible at the URL https://merenlab.org/data/pBI143 to reproduce our findings. Bacterial cultures for host range investigations, which are listed in Supplementary Table 4, are courtesy of The Duchossois Family Institute (https://dfi.uchicago.edu/). *B. fragilis* strains with pBI143 are available upon request from the Comstock Lab collection (https://comstocklab.uchicago.edu/).

## Acknowledgements

We thank the members of the Meren Lab (https://merenlab.org) and Comstock Lab (https://comstocklab.uchicago.edu/) for helpful discussions, Jason Koval for help procuring bacterial cultures, and the Duchossois Family Institute WGS facility for sequencing constructs. We thank Melinda Bootsma for help with the qPCRs on water and sewage samples. ECF acknowledges support from the University of Chicago International Student Fellowship, and ADW acknowledges support from NIGMS R35 GM133420. Additional support for ECF came from an NIH NIDDK grant (RC2 DK122394) to EBC. Authors thank The University of Chicago Center for Data and Computing for their support. This project was funded by University of Chicago start-up funds to AME.

## Author contributions

ECF and AME conceived the study. KL developed methodology. RM, EK, and AME developed computational analysis tools. ECF, MSS, PAR, ADW and AME performed formal analyses. ECF, KL, MS and LEC conducted investigations. TM, MKY, MM and SLM provided resources. ECF, MSS, ADW and AME curated data. ECF and AME prepared the figures. ECF and AME wrote the paper with critical input from all authors. LEC and AME supervised the project. EBC and AME acquired funding.

## Ethics declarations

### Ethics approval and consent to participate

Not applicable.

### Consent for publication

Not applicable.

## Supplementary Tables (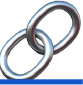)

**Supplementary Table 1:** The accompanying metadata for all publicly available metagenomes used in this study. This table contains 3 tabs. **(1)** healthy_gut: all healthy gut metagenomes. **(2)** IBD: all IBD gut metagenomes. **(3)** alternative_environment: all non-gut metagenomes.

**Supplementary Table 2:** The nucleotide sequence and average nucleotide identity (ANI) calculations for all pBI143 contigs. This table has 3 tabs. **(1)** pBI143_sequences: the nucleotide sequence for the 3 reference versions of pBI143 assembled from metagenomes. **(2)** ANI information for the 3 reference sequences of pBI143. **(3)** additional_genes: the nucleotide sequences for pBI143 assembled from metagenomes with additional genetic material.

**Supplementary Table 3:** Read recruitment data from metagenomes used in this study. This table has 4 tabs. **(1)** global adult gut metagenomes: coverage and detection data for the reference versions of pBI143 in global adult gut metagenomes **(2)** mother-infant metagenomes: coverage and detection data for the reference versions of pBI143 in mother and infant metagenomes **(3)** crassphage_comparison: coverage and detection data for crassphage in global adult gut metagenomes. **(4)** alternative_environments: coverage and detection data for the reference versions of pBI143 in non-human gut environments.

**Supplementary Table 4:** The metadata for the Duchossois Family Institute bacterial isolate genomes used in this study.

**Supplementary Table 5:** pBI143 copy number determination via qPCR. This table includes 2 tabs. **(1)** Seq DataSource: contains the Gen-Bank accession numbers and other data sources used in primer and probe development. **(2)** Hsp BLAST result: contains BLASTN results of the hsp nucleotide sequence against the 15 *Bacteroides fragilis* RefSeq complete genomes.

**Supplementary Table 6:** The data for pBI143 copy number for all animal, environmental and sewage samples as measured via qPCR. This table has 3 tabs. **(1)** animal_copy_number: contains the data showing sample and copy number of pBI143 in animal fecal samples. **(2)** environmental_copy_number: contains the data showing sample and copy number of pBI143 in water samples. **(3)** sewage_copy_number: contains the data showing sample and copy number of pBI143 in sewage samples.

**Supplementary Table 7:** All the data necessary for quantifying number and type of SNV in gut and sewage metagenomes. Variability profiles are generated by anvi’o to describe the variation found across all contigs of interest; for more information see https://merenlab.org/2015/07/20/analyzing-variability/. This table has 9 tabs. **(1)** artificial_reads_var_profile: The variability profile generated following artificial short read generation and read recruitment of pBI143 Version 2 and 3 to Version 1 (see Methods). **(2)** global_mg_var_profiles: The variability profile generated following read recruitment of all global gut metagenomes to pBI143 Version 1. **(3)** sewage_mg_var_profiles: The variability profile generated following read recruitment of all global sewage metagenomes to pBI143. **(4)** matching_SNVs_gut: The number of SNVs that do or do not match one of the reference versions of pBI143 in global gut metagenomes. **(5)** matching_SNVs_sewage: The number of SNVs that do or do not match one of the reference versions of pBI143 in global sewage metagenomes. **(6)** gut_var_profile_quince_mode: This file does not fit in excel. Link to online data to regenerate single nucleotide variant data at every position of pBI143 across all global gut metagenomes (for more details on ‘quince-mod’ see https://merenlab.org/2015/07/20/analyzing-variability/#parameters-to-refine-the-output). **(7)** sewage_var_profile_quince_mode: single nucleotide variant data at every position of pBI143 across all global sewage metagenomes. **(8)** gut_non-consensus_SNVs: Data about the plasmid version and number of non-consensus SNVs in each global gut metagenome. **(9)** sewage_non-consensus_SNVs: Data about the plasmid version and number of non-consensus SNVs in each global sewage metagenome.

**Supplementary Table 8:** This table contains SNV variability profiles for visualizing SAAVs on the pBI143 AF structure. This table has 3 tabs: **(1)** merged_variability: contains all SNV variability data calculated with ‘anvi-gen-variability-profile --engine AA’ which summarized metagenomic read recruitment results to pBI143; **(2)** merged_variability_filtered: filtered version of merged_variability that reflects the SAAV data visualized on the pBI143 structure in Fig. 3D; (3) most_prevelent_SAAVs: this tab contains a list of all SAAVs and their residue positions that are prevalent in at least 5% of samples.

**Supplementary Table 9:** The necessary data to generate pBI143 and isolate genome phylogenies. This table has 5 tabs. **(1)** amino_acid_repA_mobA_concat: the concatenated MobA and RepA sequences from all 82 isolate genomes. Concatenated genes are separated by ‘XXX’. **(2)** repA_mobA_treefile: the treefile generated from the concatenated mobA and repA sequences. **(3)** amino_acid_SCG_concat: the concatenated ribosomal protein sequences from all 82 isolate genomes. Concatenated genes are separated by ‘XXX’. **(4)** SCG_treefile: the treefile generated from the concatenated ribosomal protein sequences. **(5)** species_donor_information: the associated isolate data that matches the donor, species and pBI143 version.

**Supplementary Table 10:** The data necessary for generating and quantifying the mother-infant network based on single nucleotide variants. This table has 8 tabs. **(1)** Finalnd_variability_profile: data for all single nucleotide variants (SNVs) present in pBI143 in Finnish mother and infant metagenomes. **(2)** Sweden_variability_profile: data for all single nucleotide variants present in pBI143 in Swedish mother and infant metagenomes. **(3)** Italy_variability_profile: data for all single nucleotide variants present in pBI143 in Italian mother and infant metagenomes. **(4)** USA_variability_profile: data for all single nucleotide variants present in pBI143 in American mother and infant metagenomes. **(5)** network_data: data used to generate the network. **(6)** distance_matrix_cosine: distance matrix calculated from network data used for quantification of distances between samples. **(7)** subtracted_distance_df: quantified distance between mother and infant samples based on cosine distance matrix. **(8)** summary of pBI143 version maintenance in infants over the sampling period.

**Supplementary Table 11:** Kraken data. This table contains 2 tabs. **(1)** Kraken data for all *Bacteroides* (this includes *Phocaeicola* with old *Bacteroides* genus names) and *Parabacteroides* taxa in global gut metagenomes and the corresponding pBI143 coverage in each of these metagenomes. **(2)** Kraken data for the number of reads recruited to a *B. fragilis* compared to the coverage of pBI143 in the same metagenomes.

**Supplementary Table 12**: This table contains 2 tabs. **(1)** The data for the mouse competition experiments. **(2)** The pBI143 maintenance in culture data.

**Supplementary Table 13:** The data for pBI143 copy number for each timepoint and condition of the *Bacteroides fragilis* stress experiments in culture as measured via qPCR. This table has 2 tabs. **(1)** 214_oxidative_stress_qPCR_data: contains data on the copy number for each test condition for the *Bacteroides fragilis* 214 strain. **(2)** R16_oxidative_stress_qPCR_data: contains data on the copy number for each test condition for the *Bacteroides fragilis* R16 strain.

**Supplementary Table 14:** The calculated ACNR and necessary data for these calculations. This table has 4 tabs. **(1)** Coverage_ratio_data: the final ACNR for all predicted single hosts of pBI143 in metagenomes. **(2)** pBI143_healthy: the coverage of pBI143 in healthy gut metagenomes. **(3)** pBI143_IBD: the coverage of pBI143 in IBD gut metagenomes. **(4)** SCG_taxonomy: Link to files containing SCG coverage data.

**Supplementary Table 15:** The names and sequences of all primers and probes used in this study.

## Supplementary Figures

**Supplementary Fig. 1.**
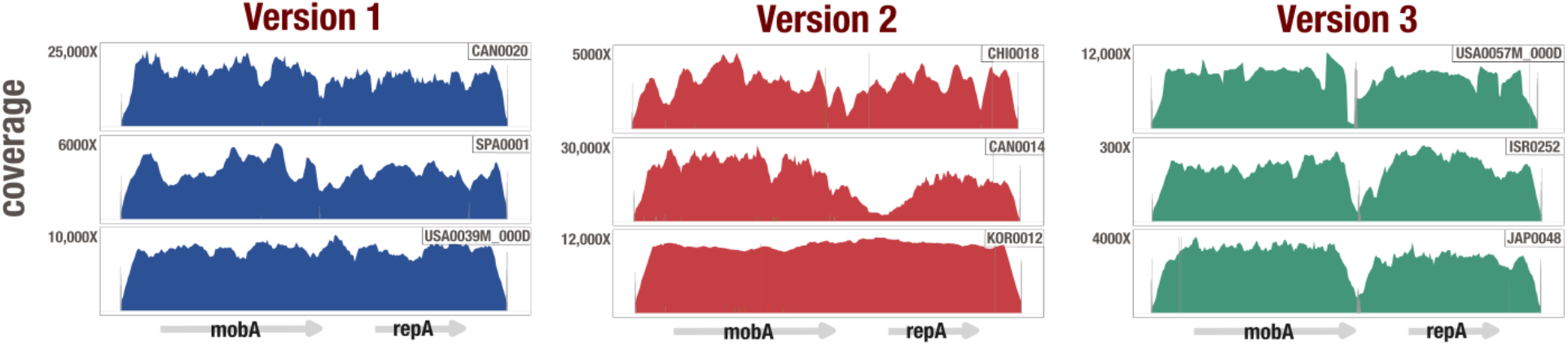
Representative coverage plots of global metagenomes mapped to pBI143. Each coverage plot shows the read recruitment results for an individual metagenome to a pBI143 Version 1 (blue), Version 2 (red) and Version 3 (green). Vertical bars show single nucleotide variants (red bar = variant in first or second codon position, green bar = variant in third codon position, gray bar = intergenic variant). The x-axis is the pBI143 reference sequence. 3 coverage plots for each reference version of pBI143 are shown, the remaining 13,539 can be generated from the anvi’o databases at https://merenlab.org/data/pBI143.

**Supplementary Fig. 2.**
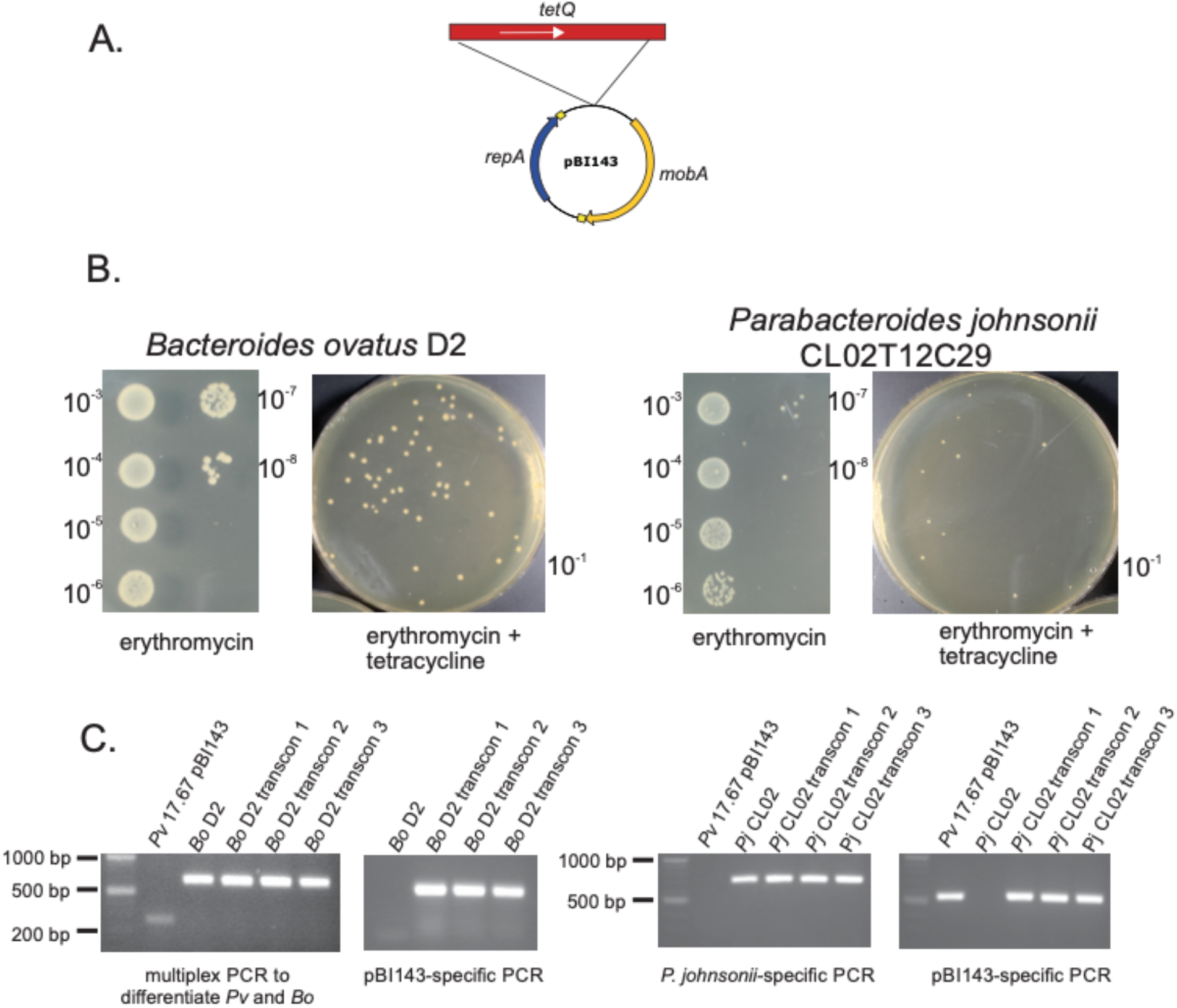
pBI143 transfer to other Bacteroidales species. **(A)** Construct made to select for plasmid transfer. **(B)** Number of recipients (erythromycin) and number of transconjugants (erythromycin and tetracycline) for transfer of pBI143-tetQ to *Bacteroides ovatus* D2 and *Parabacteroides johnsonii* CL02T12C129. **(C)** PCR to confirm presence of pBI143-tetQ in recipient strain.

**Supplementary Fig. 3.**
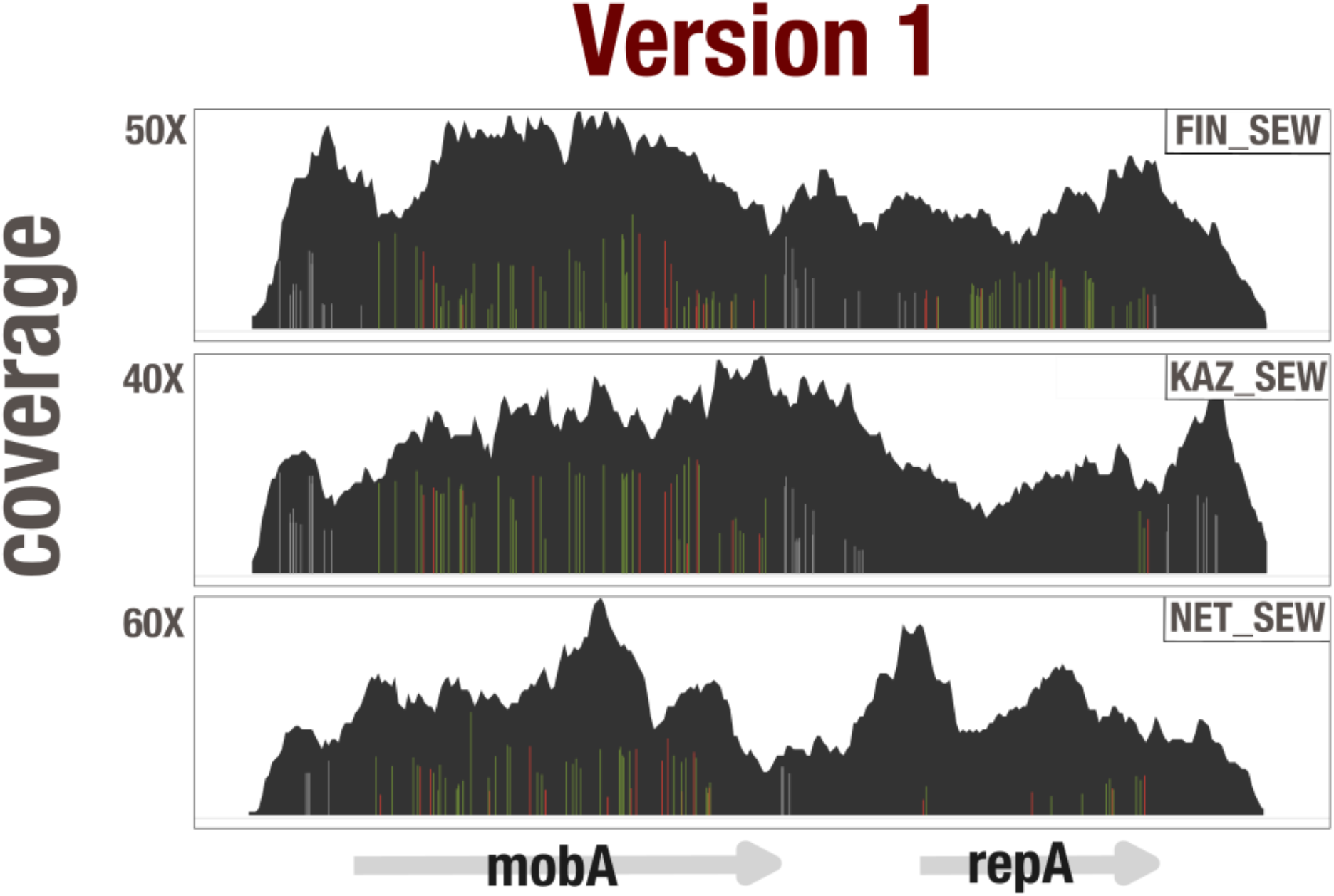
Representative coverage plots of sewage metagenomes mapped to pBI143. Each coverage plot shows the read recruitment results for a sewage metagenome to the Version 1 pBI143 reference sequence. Vertical bars show single nucleotide variants (red bar = variant in first or second codon position, green bar = variant in third codon position, gray bar = intergenic variant). The x-axis is the pBI143 reference sequence. 3 sewage coverage plots are shown, the other 435 coverage plots from all non-human environments can be generated from the anvi’o databases at https://merenlab.org/data/pBI143.

**Supplementary Fig. 4.**
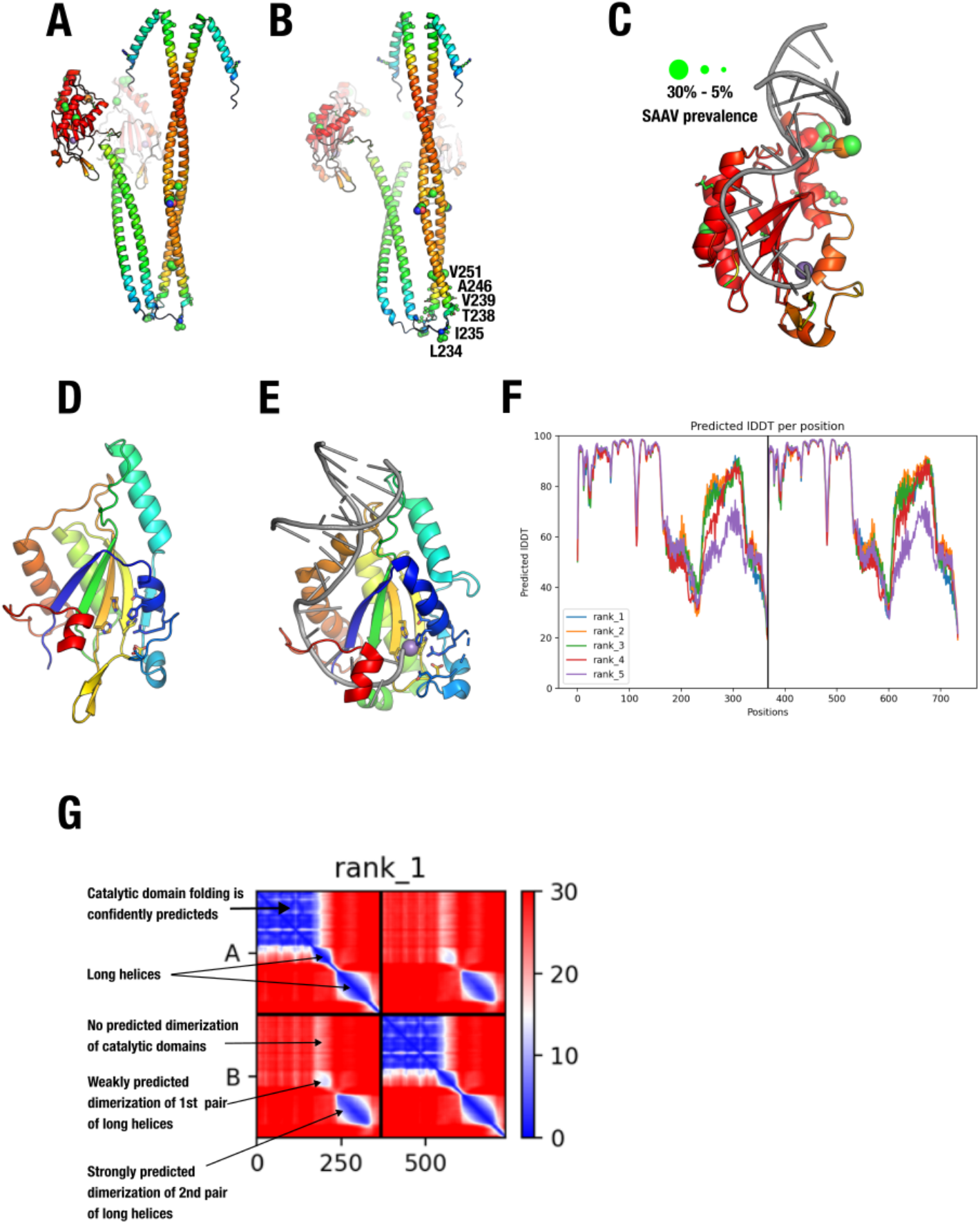
Insights from conserved single amino acid variants. **(A),(B)** Different angles of the MobA AlphaFold 2 dimer prediction with single amino acid variants from all 4,516 human gut metagenomes superimposed as ball-and-stick residues. The size of the ball-and-stick spheres indicate the proportion of samples carrying variation in that position (the larger the sphere, the more prevalent the variation at the residue) and the color is in CPK format. The color of the ribbon diagram indicates the pLDDT from AlphaFold 2 (red > 90 pLDDT) and blue < 50 pLDDT). The purple sphere is the Mn++ ion that marks the protein active site (oriT DNA and Mn2+ from 4lvi.pdb; 10.1073/pnas.1702971114). **(C)** Catalytic domain with high pLDDT with single amino acid variants from all 4,516 human gut metagenomes superimposed as ball-and-stick residues. Size and coloring is the same as in A,B. **(D)** The catalytic domain of the AlphaFold 2 predicted MobA (residues 1-177) shown shaded from blue to red active site residues are shown as sticks. **(E)** MobM from pMV158 bound to oriT DNA (gray) and a catalytic Mn2+ ion (purple) (PDB id 4lvi ^81^) shown shaded from blue to red and active site residues are shown as sticks. **(F)** AlphaFold 2 pLDDT score representing structural prediction accuracy of MobA. **(G)** AlphaFold 2 predicted aligned error plot (PAE) for MobA dimer prediction.

**Supplementary Fig. 5.**
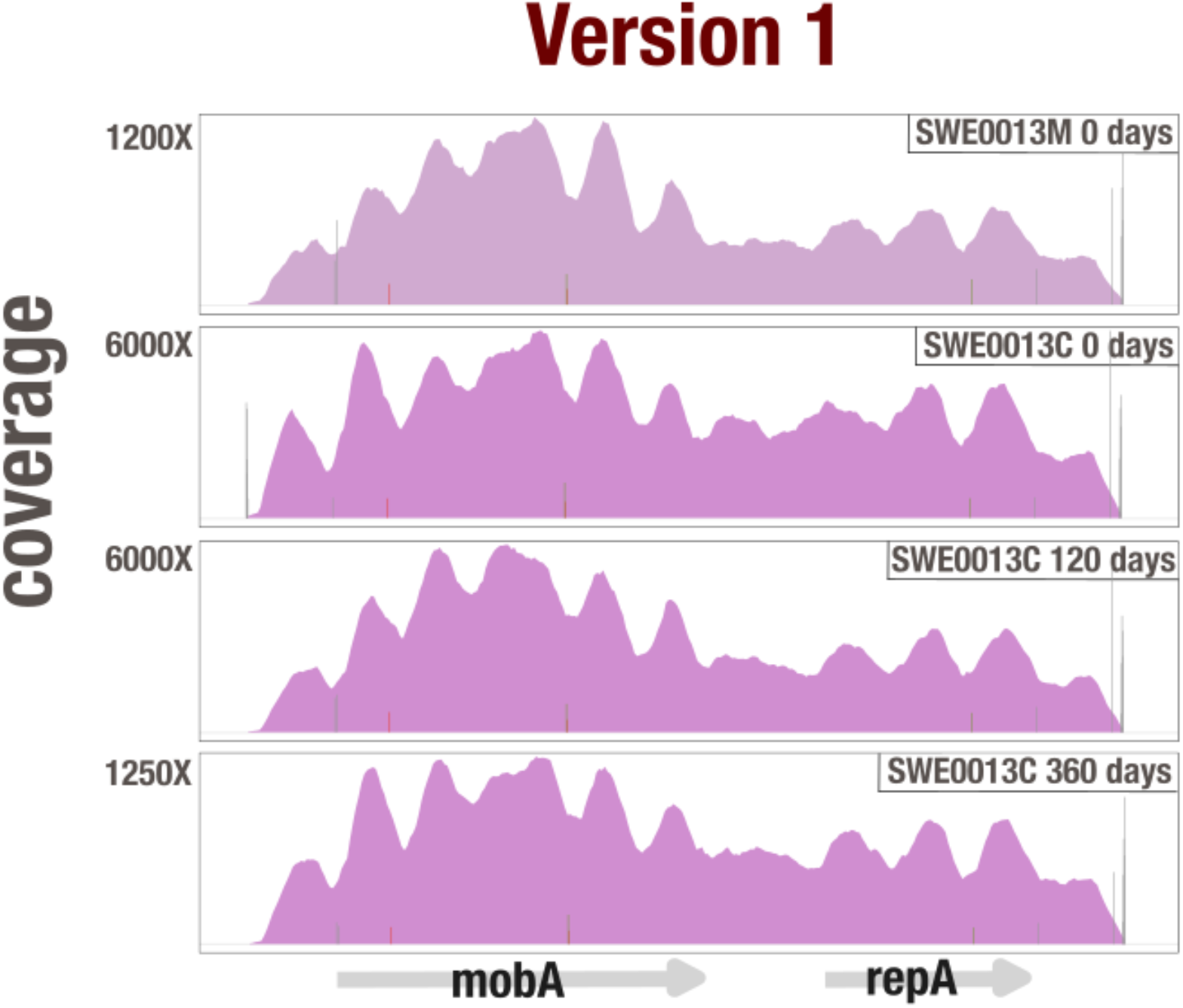
Representative mother-infant coverage plots. Each coverage plot shows the read recruitment results for an individual metagenome to the Version 1 pBI143 reference sequence. Vertical bars show single nucleotide variants (red bar = variant in first or second codon position, green bar = variant in third codon position, gray bar = intergenic variant). The x-axis is the pBI143 reference sequence. 4 coverage plots are shown, the other 1,020 can be generated from the anvi’o databases at https://merenlab.org/data/pBI143.

**Supplementary Fig. 6.**
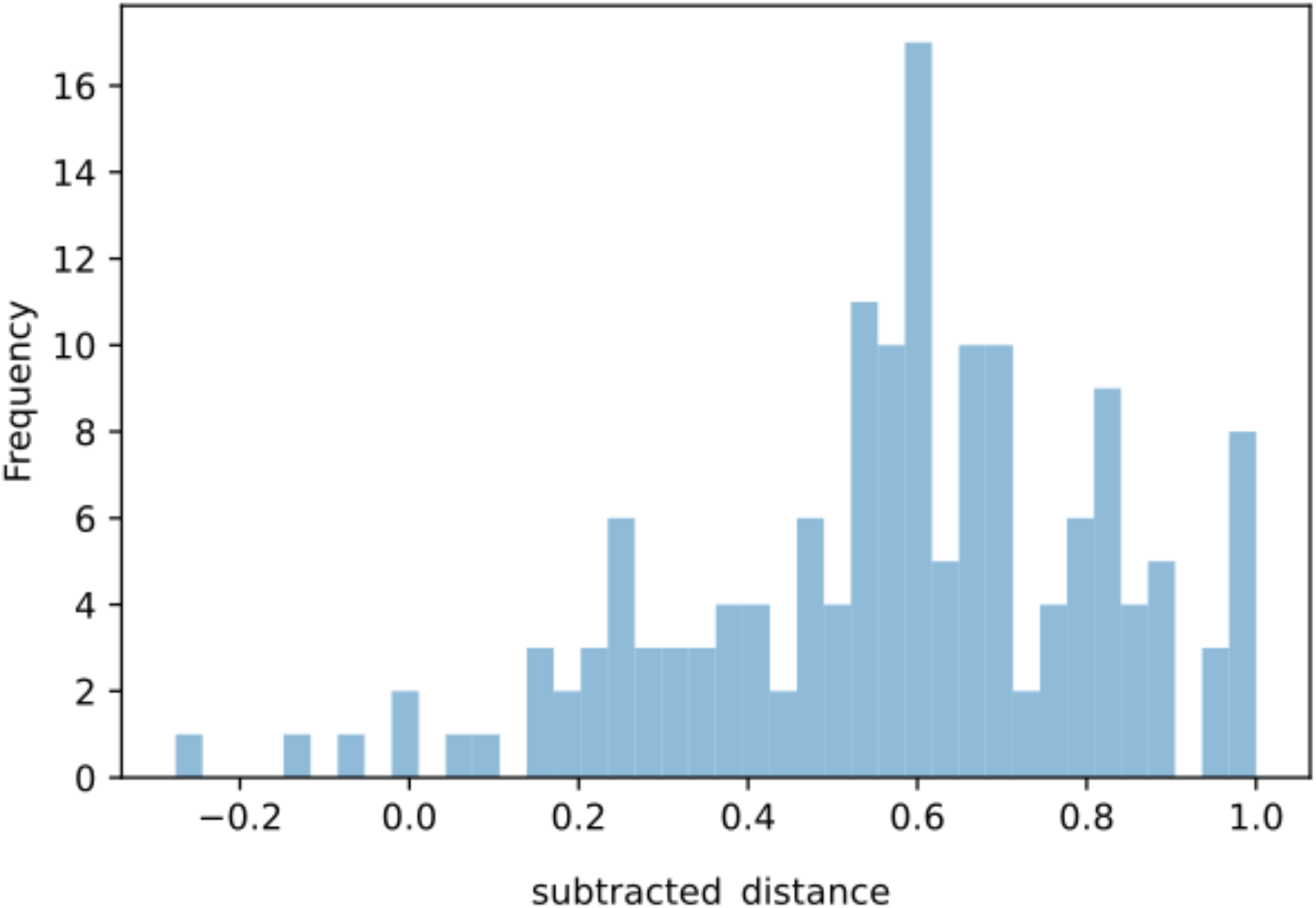
Mother-infant network quantification. Quantification of distances between samples in the network, where distance is calculated by converting the network file to a distance matrix using the python ‘pdist’function with cosine distances. The “subtracted difference” shows the mean within-family distances subtracted from mean between-family distances for each sample in the mother infant pair network. See methods for more details.

**Supplementary Fig. 7.**
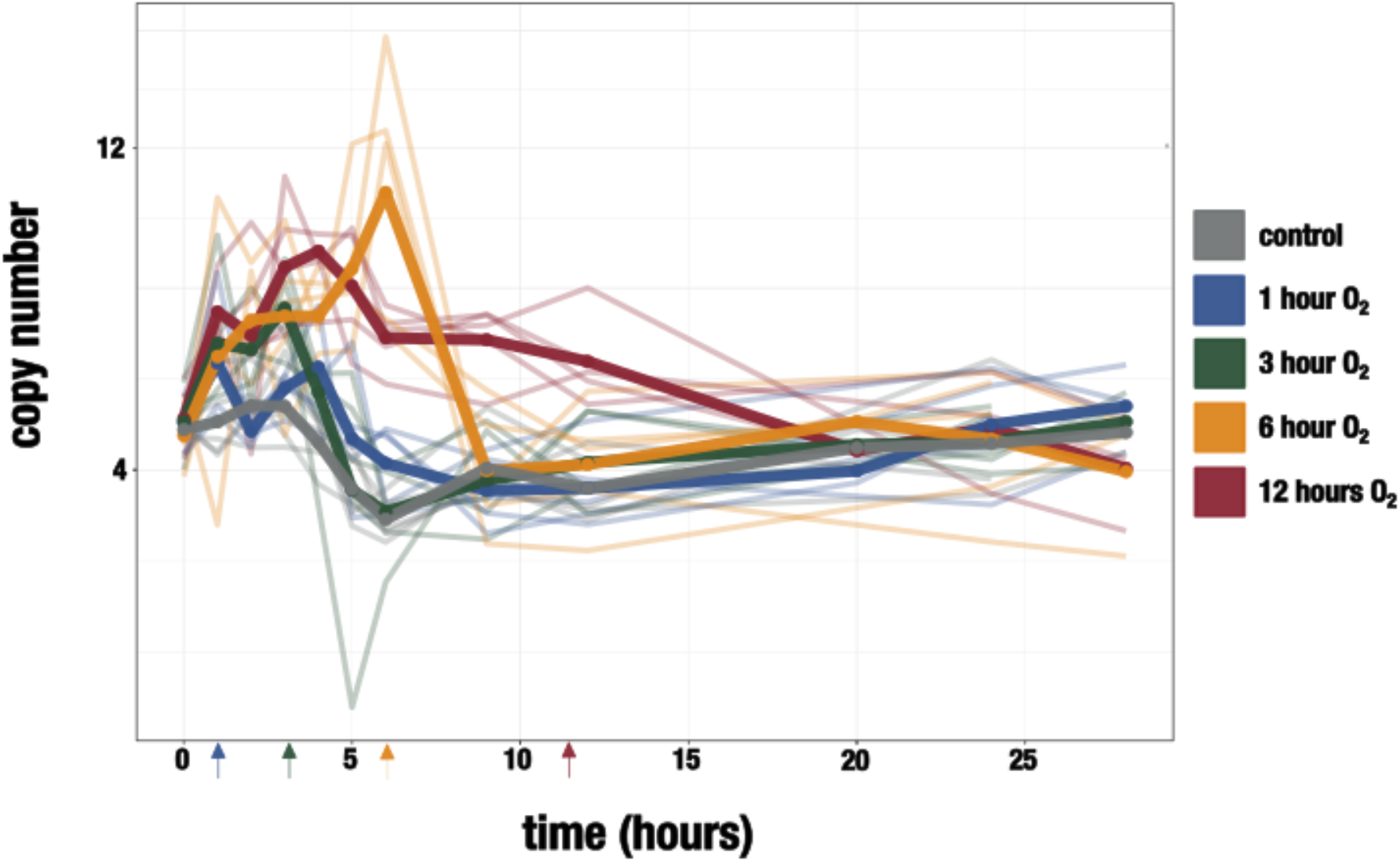
R16 oxidative stress experiment. Copy number of pBI143 in *B. fragilis* R16 cultures with increasing exposure to oxygen. Arrows indicate the time point at which the culture was returned to the anaerobic chamber. The control cultures (gray) were never exposed to oxygen. Opaque lines are the mean of 5 replicates (translucent lines).

**Supplementary Fig. 8.**
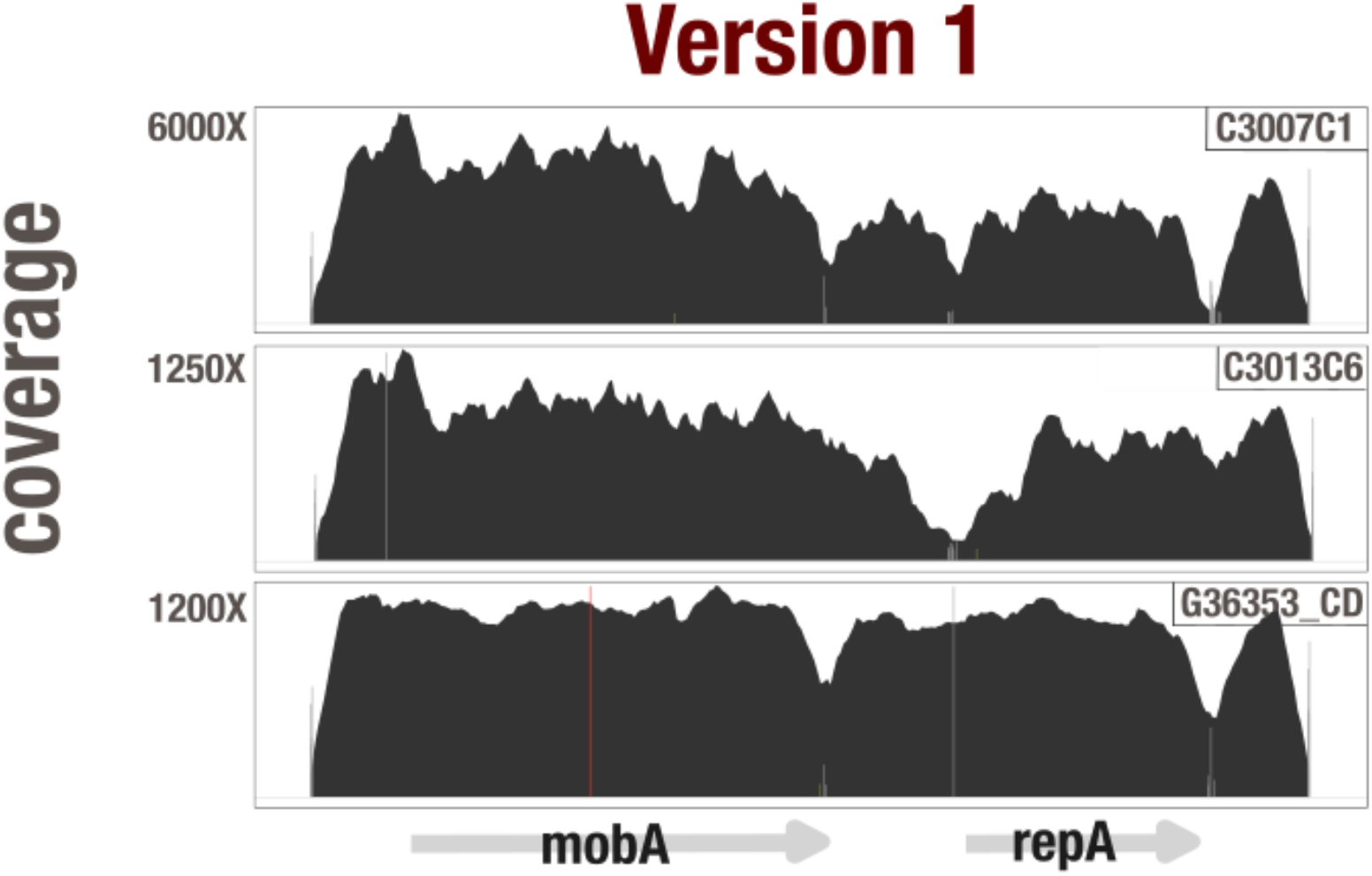
Representative IBD gut metagenome coverage plots. Each coverage plot shows the read recruitment results for an individual metagenome to a pBI143 Version 1. Vertical bars show single nucleotide variants (red bar = variant in first or second codon position, green bar = variant in third codon position, gray bar = intergenic variant). The x-axis is the pBI143 reference sequence. 3 coverage plots are shown, the other 3,087 can be generated from the anvi’o databases at https://merenlab.org/data/pBI143.

## Supplementary Information

The supplementary information file (available at <u>10.6084/m9.figshare.22336666</u>) includes additional information regarding the construction of the phylogenetic trees and plasmid constructs, the development of the qPCR assay for determining copy number of pBI143, and additional information about the read recruitment results of non-human gut environments to pBI143.

